# Interpretable brain decoding from sensations to cognition to action: graph neural networks reveal the representational hierarchy of human cognition

**DOI:** 10.1101/2022.09.30.510241

**Authors:** Yu Zhang, Lingzhong Fan, Tianzi Jiang, Alain Dagher, Pierre Bellec

**Author notes:** **Corresponding Author: Yu Zhang**, Artificial Intelligence Research Institute, Zhejiang Lab, Zhongtai Street, Yuhang District, Hangzhou 311100, Zhejiang, China, **Pierre Bellec**, Department of Psychology, University de Montreal, 4565, Chemin Queen-Mary, Montreal (Quebec) H3W 1W5.

## Abstract

Inter-subject modeling of cognitive processes has been a challenging task due to large individual variability in brain structure and function. Graph neural networks (GNNs) provide a potential way to project subject-specific neural responses onto a common representational space by effectively combining local and distributed brain activity through connectome-based constraints. Here we provide in-depth interpretations of biologically-constrained GNNs (BGNNs) that reach state-of-the-art performance in several decoding tasks and reveal inter-subject aligned neural representations underpinning cognitive processes. Specifically, the model not only segregates brain responses at different stages of cognitive tasks, e.g. motor preparation and motor execution, but also uncovers functional gradients in neural representations, e.g. a gradual progression of visual working memory (VWM) from sensory processing to cognitive control and towards behavioral abstraction. Moreover, the multilevel representations of VWM exhibit better inter-subject alignment in brain responses, higher decoding of cognitive states, and strong phenotypic and genetic correlations with individual behavioral performance. Our work demonstrates that biologically constrained deep-learning models have the potential towards both cognitive and biological fidelity in cognitive modeling, and open new avenues to interpretable functional gradients of brain cognition in a wide range of cognitive neuroscience questions.

**Highlights:**

- BGNN improves inter-subject alignment in task-evoked responses and promotes brain decoding
- BGNN captures functional gradients of brain cognition, transforming from sensory processing to cognition to representational abstraction.
- BGNNs with diffusion or functional connectome constraints better predict human behaviors compared to other graph architectures

**Graphic Abstract:** 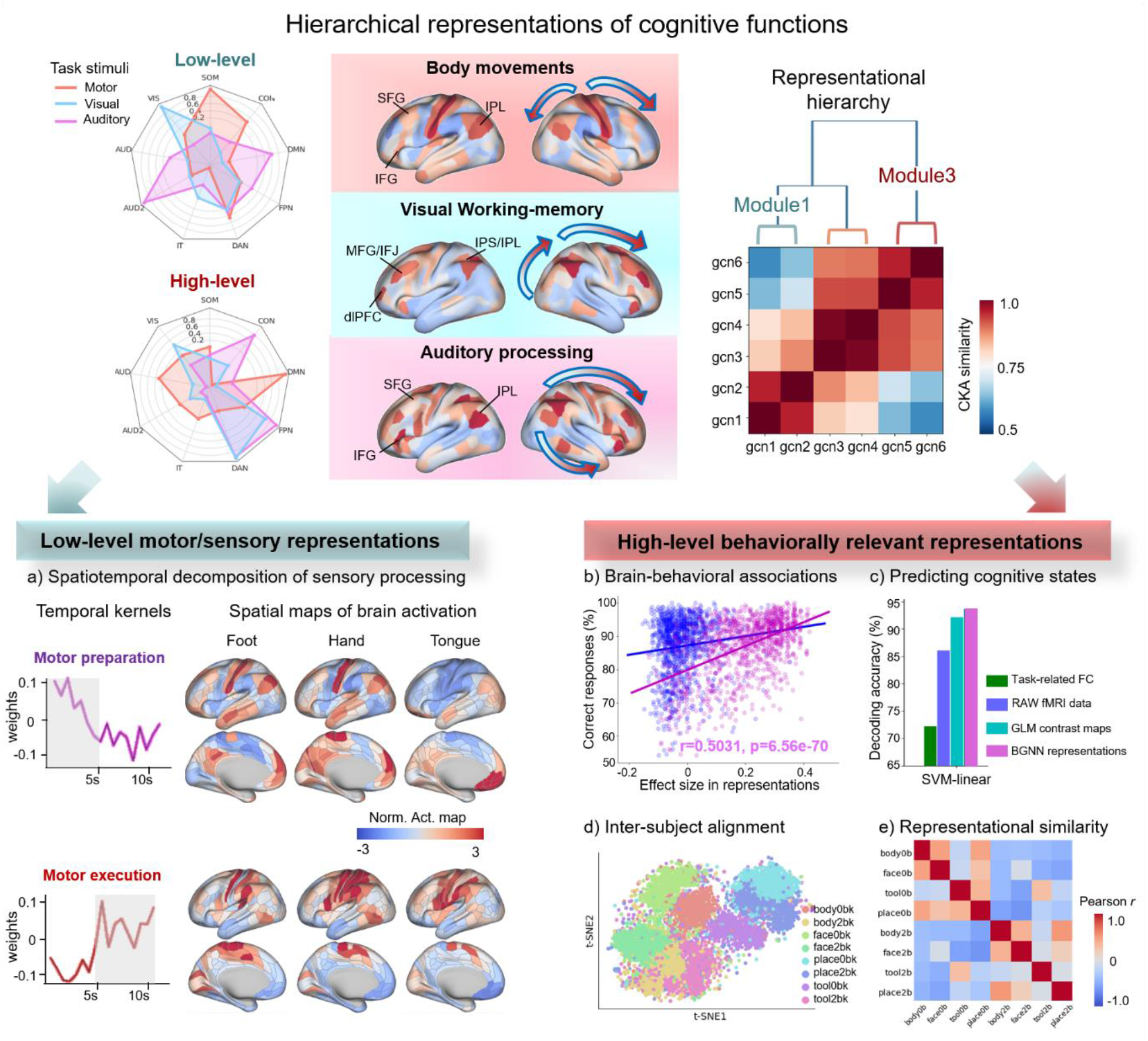

Multilevel representational learning of cognitive processes using BGNN

## Introduction

Understanding the neural substrates of human cognition is a main goal of neuroscience research. Modern imaging techniques, such as functional magnetic resonance imaging (fMRI), provide an opportunity to map cognitive function in-vivo. However, due to large inter-subject variability in brain anatomy and function, as well as in behaviors (Llera et al., 2019), modeling shared information in take-evoked neural dynamics across individuals remains challenging. To address this issue, an emerging topic of hyperalignment or functional alignment has been proposed, which aims to project subject-specific neural responses into a common representational space (Bazeille et al., 2021; Guntupalli et al., 2016; Haxby et al., 2020) using either linear transformations of neural activity (Bazeille et al., 2021; Guntupalli et al., 2016; Haxby et al., 2011) or connectivity profiles (Guntupalli et al., 2018; Levakov et al., 2021; Wang et al., 2015). Few attempts have been reported to combine both aspects of neural activity and connectivity information. As a generalization of convolutions onto high-dimensional or non-Euclidean data, graph neural networks (GNNs) provide a potential solution to integrate local and distributed brain activity through connectome-based constraints, paying the way towards the precision functional mapping of individual brains.

The majority of functional mapping approaches relied on brain activity from a local area by associating cognitive functions with different patterns of brain activation. This set of techniques have gained many successes when tackling unimodal cognitive functions, including visual features (Haxby et al., 2014, 2011; Huth et al., 2012; Naselaris et al., 2015; Nishimoto et al., 2011; Stansbury et al., 2013), auditory (Kell et al., 2018; Norman-Haignere et al., 2015) and linguistic information (Mitchell et al., 2008; Nishida and Nishimoto, 2018). Accumulated evidence strongly suggests that brain cognition requires functional integration of neural activity at multiple scales, ranging from cortical neurons to brain areas towards large-scale brain networks (Christophel et al., 2017; Pulvermüller et al., 2021). One typical example is the visual working memory task (VWM), which involves largely distributed brain networks and multilevel interactions among memory, attention and other sensory processes (Brincat et al., 2018; Christophel et al., 2017; Eriksson et al., 2015; Tang et al., 2019). For instance, the visual cortex encodes low-level sensory features, e.g., orientation (Harrison and Tong, 2009), motion (Riggall and Postle, 2012) and patterns of the visual stimuli (Christophel et al., 2012), while the prefrontal and parietal cortex maintenance the abstract representations over a delayed interval in the memory system (Christophel et al., 2012; Oh et al., 2019; Sligte et al., 2013). Studies have uncovered a gradual progression of WM from the low-level sensory processing in sensory cortices to behaviorally relevant abstract representations in prefrontal regions by using recordings of neural activity in primates (Brincat et al., 2018; D’Esposito and Postle, 2015). Accurately mapping such multilevel integrative processes of WM in the human brain is still challenging, mainly due to the high computational complexity of the full-brain models in conventional neuroimaging analysis (Haxby et al., 2014; Huth et al., 2012; Nakai and Nishimoto, 2020; Nishimoto et al., 2011) and poor inter-subject alignment of brain responses in large-scale neuroimaging data (Haxby et al., 2020; Poldrack et al., 2009).

Recently, GNNs have reached state-of-the-art performance in several brain decoding benchmarks (Hou et al., 2020; Li et al., 2021; Lin et al., 2021; Zhang et al., 2021; Zhang and Bellec, 2020), including our previous work on Human Connectome Project (HCP) tasks (Zhang et al., 2022, 2021). Our findings have demonstrated a remarkable boost in inter-subject decoding by using GNNs, as well as their ability to capture state-specific brain signatures in the spatiotemporal neural dynamics. However, the interpretability of GNNs and other deep learning models is a big challenge for cognitive modeling (Kriegeskorte and Douglas, 2018; Thomas et al., 2021). Specifically, it is still unknown why GNNs outperform the conventional univariate (Huth et al., 2012; Naselaris et al., 2015; Nishimoto et al., 2011) and multivariate analysis (Haxby, 2012; Haxby et al., 2014) in these tasks. We hypothesized that GNNs efficiently combine local and distributed brain activity through biologically constrained mechanisms (Pulvermüller et al., 2021), e.g. leveraging the inductive bias of empirical brain connectomes (Zhang et al., 2022). To test this hypothesis, we interpreted the latent space of GNN decoding models using modern feature/layer visualization techniques (Nguyen et al., 2019; Shi et al., 2020) as well as the well-established representational similarity analysis (Groen et al., 2018; Kornblith et al., 2019; Xu and Vaziri-Pashkam, 2021). The latent representations of GNN models were then mapped onto the human brain in a hierarchical manner and their biological basis were specifically investigated in terms of the correspondence with conventional univariate activation maps and the association with human behaviors and genetics.

In the current study, we propose a biologically-constrained spatiotemporal GNN architecture to encode the distributed, integrative processes of cognitive tasks and to decode task-related brain dynamics at fine timescales. We evaluate the model on the HCP task-fMRI database consisting of 1200 healthy subjects (Van Essen et al., 2013) and investigate the reliability and interpretability of the latent representations on a variety of cognitive tasks, including motor and perception as well as high-order cognitive functions. Taking Motor and WM tasks as examples, we systematically investigate the interpretability of the connectome-constrained GNNs, including 1) multilevel representational learning of cognitive processes, transforming from low-level sensory processing to high-level behaviorally relevant abstract representations following the cortical hierarchy; 2) spatiotemporal decomposition of cognitive tasks into multiple temporal stages and activating different brain systems; 3) inter-subject alignment of task-related neural responses and their associations with cognitive behaviors and genetic variances; 4) salient state-specific neuroimaging features and their inter-trial/subject stability. The present study provides a novel perspective of interpreting GNN models for large-scale cognitive decoding and highlights three core components for cognitive modeling, i.e. brain connectome, functional integration and representational hierarchy, which might be the keys towards brain-inspired artificial intelligence of human cognition.

## Results

### Summary of the main results

Our BGNN encoding-decoding model of cognitive functions (as shown in Fig. **1**) learns multilevel latent representations transforming from sensory processing to representational abstraction (*encoding* phase) and predicts cognitive states using embedded representations at fine timescales (*decoding* phase). First, the embedding model (Fig. **1a**) projects the high-dimensional task-evoked whole-brain activity into a dynamic brain graph and learns embedded representations through multi-layer graph neural networks. Second, the encoding model (Fig. **1b**) reveals a representational hierarchy underpinning cognitive processes, e.g. a functional gradient in neural representations of visual working memory (VWM) from low-level motor/sensory inputs to high-level abstract representations. At the low-level representations, the model uncovers spatiotemporal decompositions of task-related brain responses, i.e. decomposing cognitive processes into multiple temporal stages (e.g. *motor execution* and *motor preparation* for Motor tasks) and capturing different patterns of spatial activation maps at each stage (e.g. prefrontal regions for *motor preparation* and sensorimotor cortices for *motor execution*). At the high-level representations, the model learns behaviorally relevant abstract representations of cognitive functions that further associate with participants’ in-scanner task performance (e.g. correct responses and response time of WM tasks) and improve the inter-subject alignment of brain responses. Using the high-level representations, the decoding model (Fig. **1c**) achieves state-of-art decoding performance on a variety of cognitive functions at multiple timescales (Table 2 and Fig.6-S2), for instance, on unimodal cognitive functions like Language (F1-score = 98.36%, 2 conditions, story vs math) and Motor tasks (98.01%, 5 conditions, left/right hand, left/right foot and tongue), as well as high-order cognitive processes including Working-Memory tasks (94.14%, classifying 8 conditions, combination of the category recognition task and N-Back memory task). We will explain these key findings in more detail in the following sections.

**Fig. 1.**
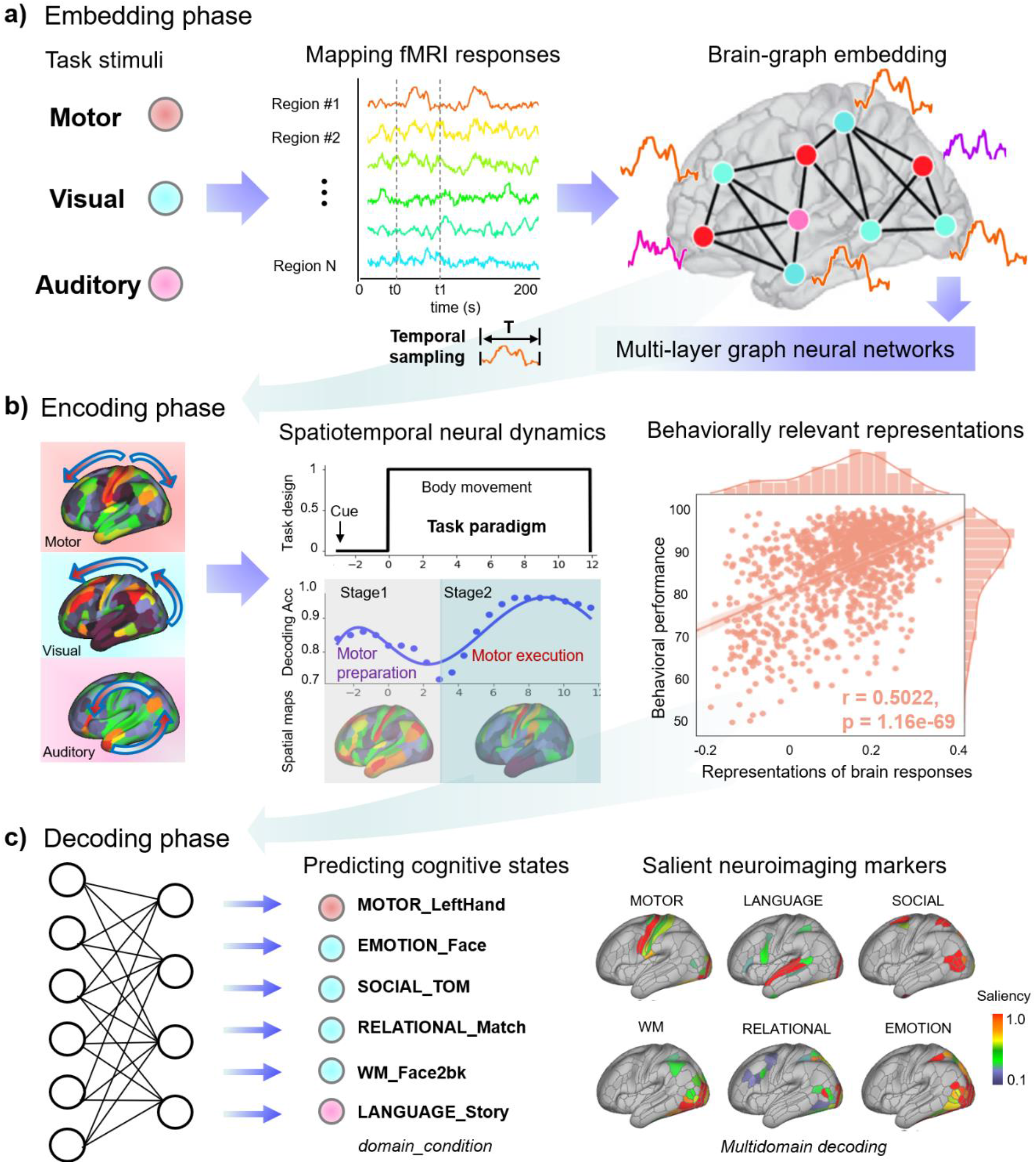
Encoding-decoding model of human cognitive functions using graph embeddings. The model consists of three stages, i.e. graph embedding, encoding and decoding. The embedding phase (**a**) maps task-related fMRI responses onto a dynamic brain graph. The encoding phase (**b**) captures hierarchical representations of cognitive functions using connectome-constrained BGNN, representing a gradual progression from motor/sensory inputs (i.e. motor/visual/auditory) to behaviorally relevant abstract representations. The decoding phase (**c**) infers cognitive states from encoded high-level BGNN representations with fine temporal resolution and fine cognitive granularity.

### Sensory-cognition-behavior representational hierarchy of WM tasks

The encoding model captures a representational hierarchy of task-related brain responses along BGNN layers. Specifically, in early BGNN layers, the model learns low-level representations of brain responses underpinning motor/visual/auditory processing, i.e. decomposing brain activity into multiple temporal stages and the corresponding spatial maps of brain activations (Fig. **4** and Fig. **4-S1**). In deep layers, the model learns high-level abstract representations of cognitive processes that are strongly associated with participants’ behavioral performance (Fig. **5** and Fig.**5**-S1). To verify this, we evaluated the representational similarity of the BGNN model using centered kernel alignment (CKA) with a linear kernel (Kornblith et al., 2019), with 0 <CKA<1, and revealed a hierarchical organization of the embedded representations for each cognitive domain using Ward linkage.

A three-level representational hierarchy was revealed for WM tasks (as shown in Fig.**2a**), including low-level features (gcn1 to gcn2), hidden representations (gcn3 to gcn4), and high-level representations (gcn5 to gcn6). Among which, early BGNN layers extracted sensory processing information in the ventral visual stream, middle BGNN layers retrieved cognitive control signals in the frontoparietal regions, and the last BGNN layer (gcn6) captured behaviorally relevant representations in the prefrontal cortex and salience network (Fig. **2d**). These BGNN representations demonstrated weak associations between different representational levels (CKA=0.94 and 0.76 for within- and between-level similarity), with a stepwise progression from sensory processing to cognitive control and towards behavioral abstraction (CKA=0.54, 0.83, 0.92 for low, middle, high-level features as compared to gcn6). Moreover, the high-level BGNN representations demonstrated a strong category-specific effect by learning similar features for the same task but showing distinct features between tasks (Fig.**2b** and **c**). This category-specific effect was gradually enhanced along the representational hierarchy (Fig. 5-S2a) and all BGNN representations demonstrated higher contrasts of 2back vs 0back tasks (*2bk-0bk*) compared to the GLM-derived contrast maps (Fig. 2-S1b and Fig. 5-S2b). The representational hierarchy of WM tasks resembled the previously reported progression of activity flow in WM tasks, i.e. information transformation from sensory inputs to behaviorally relevant representations along the cortical hierarchy, as revealed by neural recordings in macaques (Brincat et al., 2018).

**Fig. 2.**
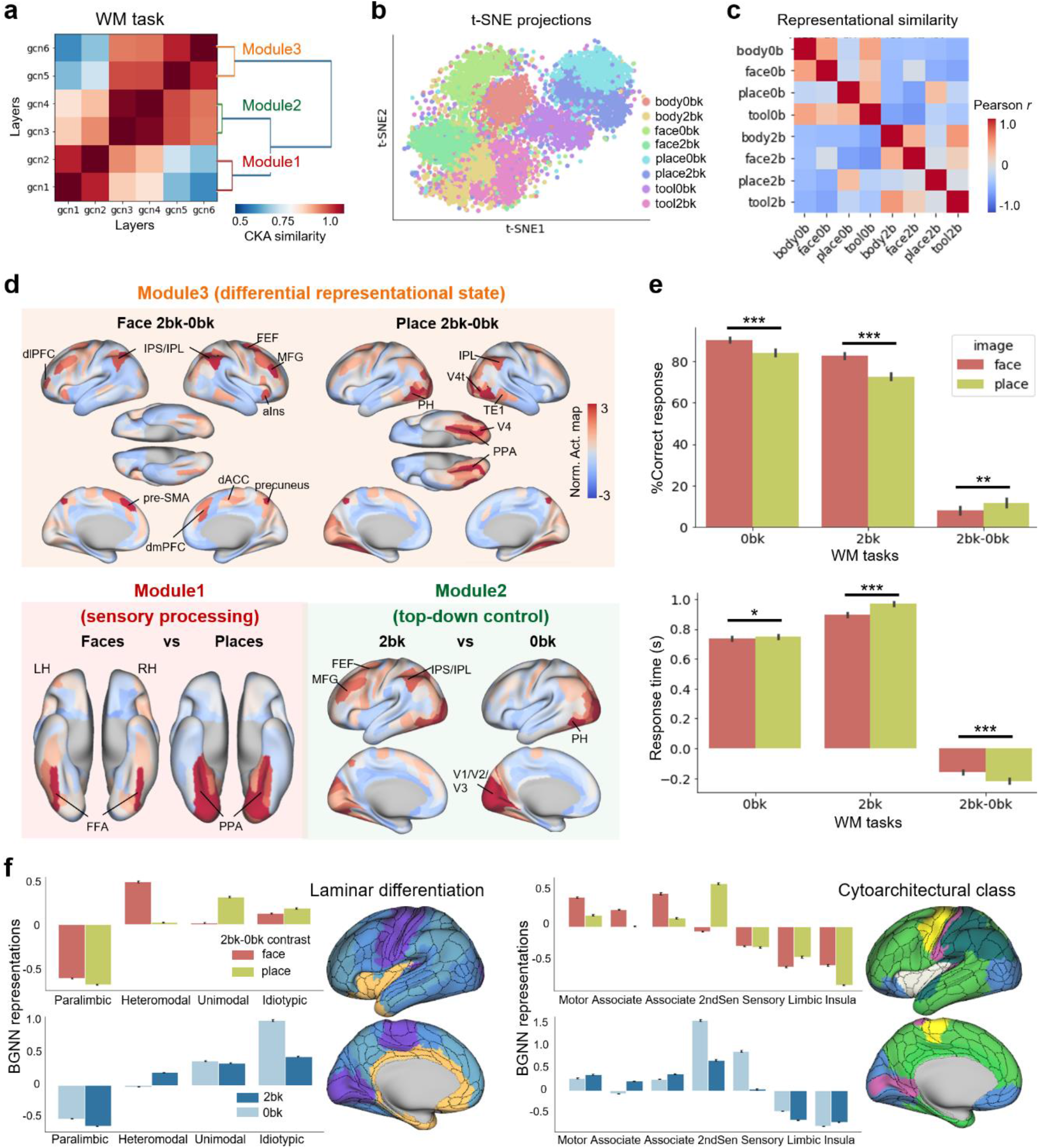
BGNN revealed a representational hierarchy of VWM tasks transforming from sensory processing in visual areas to behavioral abstraction in prefrontal cortices. **a)**, We found a three-level representational hierarchy of WM tasks by using centered kernel alignment (CKA) to evaluate the similarity of BGNN representations and performing hierarchical clustering on the similarity matrix. **b)**, Representations of WM tasks in the last BGNN layer (gcn6, part of *Module3*) exhibited a strong task-specific effect of t-SNE projections, with distinct clusters for each task condition and small overlaps between tasks. **c)**, The representational similarity, evaluated by Pearson correlation coefficients, demonstrated highly discriminative BGNN representations between 2back and 0back tasks as well as among tasks using different visual stimuli, e.g. faces vs places. **d)**, Multilevel representational learning of WM tasks. *Module1* (in the red block) detected neural representations of visual processing, e.g. the recognition of face and place images in the ventral stream. *Module2* (in the green block) detected neural representations of memory load, e.g. the contrast of *2bk vs 0bk* tasks in the frontoparietal regions. *Module3* (in the orange block) revealed divergent brain mechanisms for the *2bk-0bk* contrasts on familiar faces and places, indicating a differential representational state for recognizing familiar faces. **e)**, A privileged WM state for familiar faces in human behavioral data. Participants remembered better (i.e. higher accuracy and faster responses) on familiar faces than places for both 0back (*0bk*) and 2back tasks (*2bk*), and showing smaller decays due to the memory load (*2bk-0bk*). *** indicates p-value<0.001, ** indicates p-value<0.01, * indicates p-value<0.05. **f)**, Spatial associations between BGNN abstract representations and levels of laminar differentiation (left) and cytoarchitectural taxonomy (right). dlPFC: dorsolateral prefrontal cortex; dmPFC: dorsal medial prefrontal cortex; MFG: middle frontal gyrus; IFJ: inferior frontal junction; aIns: anterior insula; dACC: dorsal anterior cingulate cortex; pre-SMA: pre-supplementary motor area; FEF: frontal eye fields; IPS: intraparietal sulcus; IPL: inferior parietal lobule; FFA: fusiform face area; PPA: parahippocampal place area; V4t: V4 transition zone; TE1: visual processing area of the inferior temporal cortex.

Our results also revealed a 3-fold separation of neural basis underlying the information processing of WM tasks (Fig. **2d**). First, the separation of sensory processing, e.g. recognition of face *vs* place images (*face-place*), was reliably captured in the ventral stream, e.g. fusiform face area (FFA) and parahippocampal place area (PPA), in *Module1* (Fig. **2d**), consistent with the well-known segregation of the neural substrates for encoding faces and places respectively (Golarai et al., 2007). Second, the *2bk-0bk* separation was weakly detected in *Module1* but demonstrated a strong separation effect in *Module2*, especially in the frontoparietal regions including frontal eye fields (FEF), middle frontal gyrus (MFG), intraparietal sulcus (IPS) and inferior parietal lobule (IPL). These detected regions coincided with the current view of prefrontal top-down control over sensory processing in N-back tasks (Christophel et al., 2017; D’Esposito and Postle, 2015; Nee and D’Esposito, 2018). Third, the memory-vs-content disassociation was additionally captured in *Module3*, suggesting a content-specific memory mechanism. Specifically, *Module3* revealed distinct neural mechanisms underlying the contrast of *2bk-0bk* on familiar faces vs places aside from the common frontoparietal basis of *2bk-0bk* separation. The *2bk-0bk* contrast on face images relied more on top-down modulation from prefrontal cortex and salience network including the dorsolateral prefrontal cortex (dlPFC), anterior insula (aIns) and anterior cingulate cortex (ACC). By contrast, the *2bk-0bk* contrast on place images relied more on bottom-up sensory inputs in the lateral occipito-temporal cortex, including PPA, V4 and TE1.

Our findings of the memory-vs-content dissociation in both BGNN representations and neural substrates of WM tasks support the theory of a task-dependent prefrontal-vs-sensory contribution in cognitive tasks such that the sensory perception relies on sensory cortices while representational abstraction relies on prefrontal regions (Christophel et al., 2017; Nee and D’Esposito, 2018). Coincidingly, participants’ in-scanner behavioral performance also confirmed the divergent mechanisms for remembering faces and places in WM tasks and exhibited a preferential effect towards the recognition of faces. As shown in Fig.**2e**, participants better remembered familiar faces than places, by achieving higher accuracies and faster responses on both *0bk* (T=7.76, p=1.84e-14 for Acc, T=-2.38, p=0.017 for RT) and *2bk* tasks (T=12.22, p=2.86e-32 for Acc, T=-9.90, p=3.68e-22 for RT), and showing smaller decays in behavioral performance due to the increase of cognitive demands (i.e. *2bk-0bk*, T=3.21, p=0.0013 for Acc, T=-5.97, p=3.16e-9 for RT). Our findings coincided with the literature on a privileged WM state for faces with an improved accuracy and response time in both newborns and adults (Farroni et al., 2005; Lin et al., 2019; Sato and Yoshikawa, 2013). Together, both neural activity and behavioral data supported the 3-level representational hierarchy of WM tasks and suggested a differential representational state for faces compared to non-faces.

Moreover, in order to validate the biological basis of such representational hierarchy and the memory-vs-content disassociation of WM tasks, we mapped the embedded BGNN representations onto independent atlases of laminar differentiation (Mesulam, 1998) and cytoarchitectural class. We found that *2bk* tasks relied on the association cortices while the *0bk* tasks relied on the primary and secondary sensory cortices (Fig.**2f**). By contrast, we observed divergent neural substrates underlying the *2bk-0bk* contrasts, i.e. heteromodal association areas for faces and unimodal sensory areas for places (Fig.**2f**).

### Functional gradient of Motor tasks: from motor execution to motor planning

We uncovered a two-level representational hierarchy of Motor tasks (as shown in Fig.**3**), including the low-level sensory processing (gcn1 to gcn2) and high-level abstract representations (gcn3 to gcn6). Among which, we detected weak associations between two representational levels (CKA=0.58 for gcn1-gcn2 as compared to gcn6), along with highly redundant features in the hidden representations (CKA=0.92 for gcn3-gcn5 as compared to gcn6). The low-level sensory processing decomposed task-evoked brain activity in both spatial and temporal domains, as revealed by the feature visualization of spatiotemporal graph filters in the 1st BGNN layer, showing biologically relevant activation patterns in the sensorimotor and prefrontal cortices (Fig.**3c**). The high-level abstract representations captured the intention of movements and demonstrated an evident task-specific effect that showing similar features for the same type of body movements, including left and right body parts, and distinct features among different body movements (Fig.**3b**). The follow-up representational similarity analysis exhibited much higher contrasts of different body movements as compared to the classical GLM analysis (Fig.2-S1a). The representational hierarchy of Motor tasks identified two phases of motor processes, i.e. *motor planning* and *motor execution*, and uncovered a functional gradient in the neural representations of Motor tasks following the cortical hierarchy (Fig.**3d**). Specifically, the execution phase of body movements was detected in *Module1* by revealing well-established activation patterns in the sensorimotor cortex. The planning phase of movements was captured in *Module2* by involving the prefrontal and parietal regions during all Motor tasks, including medial prefrontal cortex (mPFC), inferior frontal gyrus (IFG), FEF and IPL. Our findings uncovered the spatiotemporal dynamics underlying motor processes and revealed distinct neural substrates for the stages of motor tasks, i.e. motor execution and motor planning. Our results indicated a potential role of frontoparietal regions in the planning of goal-directed actions. Similar two-stage functional segregation of motor processes has been reported in humans (Ariani et al., 2022; Gallivan et al., 2011), monkeys (Messinger et al., 2021) and rodents (Eriksson et al., 2021).

**Fig. 3.**
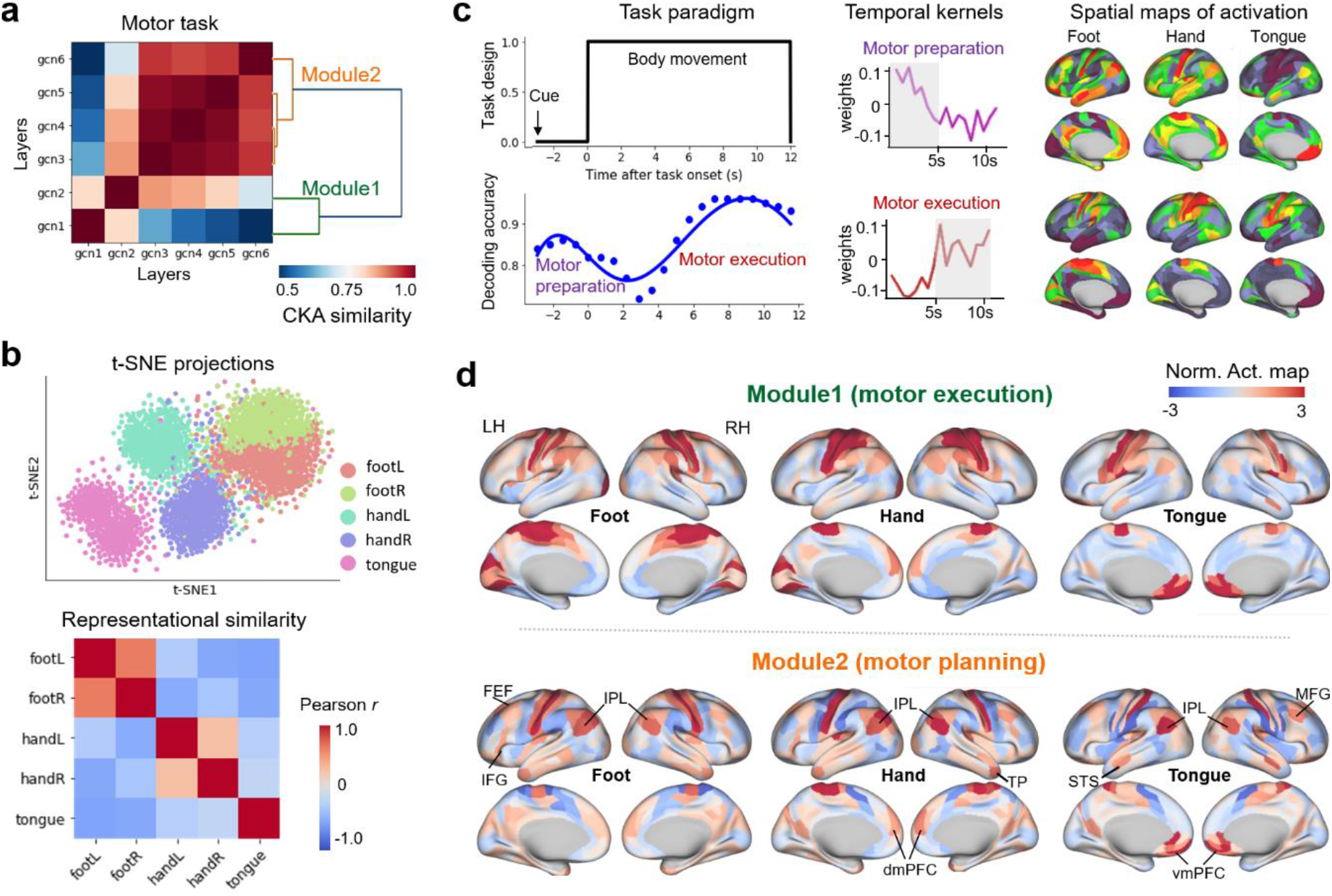
Hierarchical organization of BGNN representations for the MOTOR tasks. **a)**, We found a two-level representational hierarchy of Motor tasks by using CKA to evaluate the similarity of BGNN representations and performing hierarchical clustering on the similarity matrix. **b)**, Representations of Motor tasks in the last BGNN layer (gcn6, part of *Module2*) exhibited a strong task-specific effect in t-SNE projections. The representational similarity, evaluated by Pearson correlation coefficients, demonstrated highly discriminative BGNN representations among different movement types (foot vs hand vs tongue) as well as between left and right body parts. **c)**, Spatiotemporal decomposition of the Motor process. The single-volume prediction (1^st^ panel, 2^nd^ row in **c**) indicated two peaks in the temporal curve of decoding accuracy, corresponding to two different stages of Motor tasks, i.e. motor execution (in red) and motor preparation (in violet). BGNN uncovered different shapes of temporal kernels and distinct patterns of spatial activations for the two stages, e.g. the sensorimotor cortex for motor execution, prefrontal and parietal regions for motor preparation. **d)**, Multilevel representational learning of Motor tasks. *Module1*(in green) revealed brain activations in the motor and somatosensory cortices for the execution of movements. *Module2* (in orange) detected brain activations in the prefrontal and parietal regions, which may correspond to the intention and planning of movements. MFG: middle frontal gyrus; IFG: inferior frontal gyrus; FEF: frontal eye fields; IPL: inferior parietal lobe; dmPFC: dorsal medial prefrontal cortex; vmPFC: ventral medial prefrontal cortex; STS: superior temporal sulcus; TP: temporal pole.

### Spatiotemporal decomposition of brain responses in early BGNN layers

The encoding model learns rich representations of brain responses underlying cognitive processes, as revealed by the feature visualization of spatiotemporal graph filters in the 1st BGNN layer, to decompose the entire process into multiple temporal stages and extract the corresponding maps of brain activation at each stage. For instance, in the Motor tasks, the model captured a series of activation maps corresponding to different stages of motor processes (Fig.**3c**), e.g. the prefrontal and parietal regions were involved at the *preparation* stage, i.e. neural activity immediately after the presentation of the cue images, while the sensorimotor cortex was activated during *motor execution*. Besides, the model learned a variety of temporal convolutional kernels, corresponding to the diverse shapes of hemodynamic responses (HRF, as shown in Fig.**4**). For instance, the model learned redundant convolutional kernels for the execution stage of body movements (Fig.**4b** and **d**), accounting for the variability of HRF among trials and subjects (Aguirre et al., 1998; Neumann et al., 2003). In addition, some instantaneous subprocess of cognitive functions was also captured, e.g., the visual cortex was involved for recognizing the cue images shown in the middle of a Motor task block (Fig.**4e**). This spatiotemporal decomposition of motor processes coincided with previous studies that clustered brain responses into different stages and networks in a sequential motor task (Orban et al., 2015). Using the same procedure, we observed a rich set of spatiotemporal representations underlying the Language tasks as well, corresponding to different stages of semantic and arithmetic processes (Fig.**4-S1**), for instance, the involvement of visual cortex during the *cue phase*, the engagement of prefrontal and temporal regions at the stage of *language comprehension*, the activation of sensorimotor cortex at the stage of *button pressing*. When the time window of an entire Language trial was analyzed, corresponding to the continuous stimuli of auditory processing in the fMRI paradigm, the extracted spatial maps coincided with the activation maps derived from classical GLM analysis (Fig.**4-S1b**). We did not observe such temporal decomposition for the cognitive process of WM tasks, mainly due to the lack of a clear delayed period in the N-back fMRI paradigm which makes it hard to distinguish the maintenance and retrieval periods in a single WM trial (Pinal et al., 2014). Together, the encoded low-level sensory representations uncover a sequential gradient in the spatiotemporal organization of cognitive processes, not only to distinguish patterns of brain activation in the spatial domain but also to decompose temporal dynamics of cognitive processes into multiple stages.

**Fig. 4.**
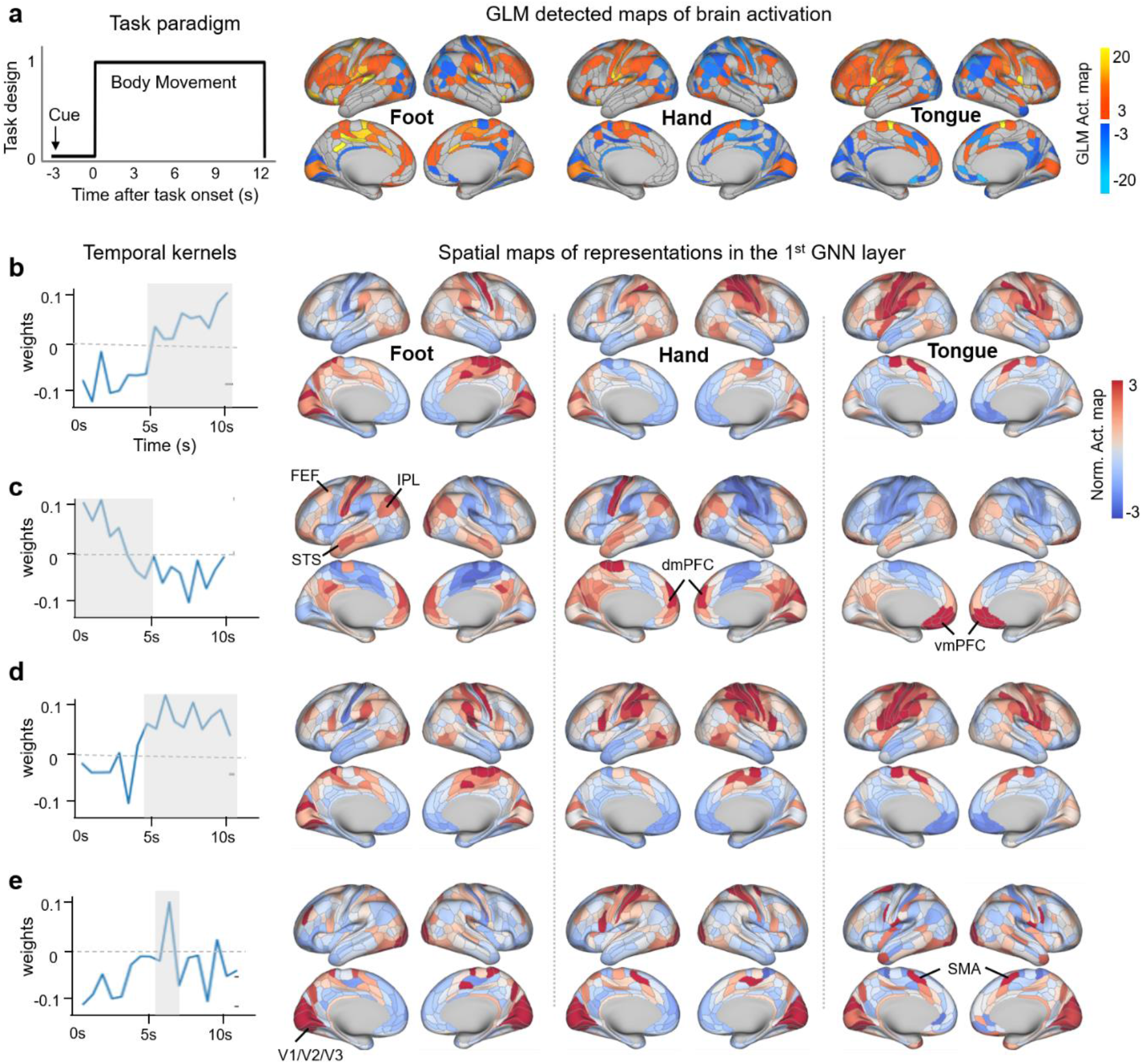
Spatiotemporal decomposition of low-level BGNN representations for Motor tasks. BGNN uncovered a multi-stage spatiotemporal organization of cognitive processes, including diverse hemodynamic responses in the temporal domain and distinct patterns of activation maps in the spatial domain. **a)**, Task paradigm of Motor trials and the corresponding activation maps detected by the classical GLM analysis. Each task block of a movement type (hand, foot or tongue) is preceded by a 3s cue and lasts for 12s. **b-e)**, BGNN captured a variety of temporal convolutional kernels (1^st^ column) corresponding to task-evoked responses at different stages of cognitive processes, for instance, the *motor preparation* (**c**) and *motor execution* (**b** and **d**), as well as processing visual cues in the middle of a task block (**e**). At each stage, the corresponding “activation maps” (2^nd^ to 4^th^ column) demonstrated distinct neural basis among task conditions, e.g. foot (2^nd^ column), hand (3^rd^ column), and tongue (4^th^ column). Our results indicated a functional gradient in the spatiotemporal organization of Motor tasks, e.g., the sensorimotor cortex for the stage of *motor execution*; prefrontal regions and default mode network (DMN) for the stage of *motor preparation*; the visual cortex for processing visual cues. FEF: frontal eye fields; IPL: inferior parietal lobe; dmPFC: dorsal medial prefrontal cortex; vmPFC: ventral medial prefrontal cortex; SMA: supplementary motor area; STS: superior temporal sulcus.

### Encoding behaviorally relevant abstract representations in deep BGNN layers

#### Improved inter-subject functional alignment of task-related brain responses

The BGNN model projects task-evoked brain responses onto a common representational space by using a graph embedding approach constrained by human connectome priors, and consequently improves the inter-subject alignment of neural responses underlying cognitive functions. Studies have shown that the inter-subject variability in brain structure and function may be a major obstacle towards a unified encoding model of cognitive processes (Bazeille et al., 2021; Haxby et al., 2020). To tackle this problem, BGNN took into account the individual variability of task-related neural dynamics at multiple scales. First, the inter-trial and inter-subject variability of HRF was embedded in early BGNN layers by learning a variety of graph convolutional kernels in the temporal domain, accounting for different stages of cognitive processes and variable shapes of HRF (Fig.**4** and Fig.**4-S1**). Second, the inter-subject variability in cognitive behaviors was encoded in deep BGNN layers by mapping subject-specific patterns of neural activity in task-related brain regions and networks (Fig. **6**) and extracting behaviorally relevant abstract representations through connectome-constrained graph convolutions (Fig. **5**). As a result, BGNN representations highly improved the functional alignment of cognitive tasks, i.e. strengthening the main effect of task conditions in neural representations while reducing between-subject variability, as compared to other commonly used neural representations, including raw fMRI data and GLM contrast maps. For instance, the representational similarity analysis demonstrated higher contrasts of different task conditions in BGNN representations than the conventional GLM contrast maps (Fig. 2-S1). An alternative dimensional reduction approach using t-SNE (Maaten and Hinton, 2008) also exhibited a stronger task-segregation effect in BGNN representations, i.e. grouping brain responses into clusters of task conditions, than raw fMRI data and GLM contrast maps (Fig.5-S3).

**Fig. 5.**
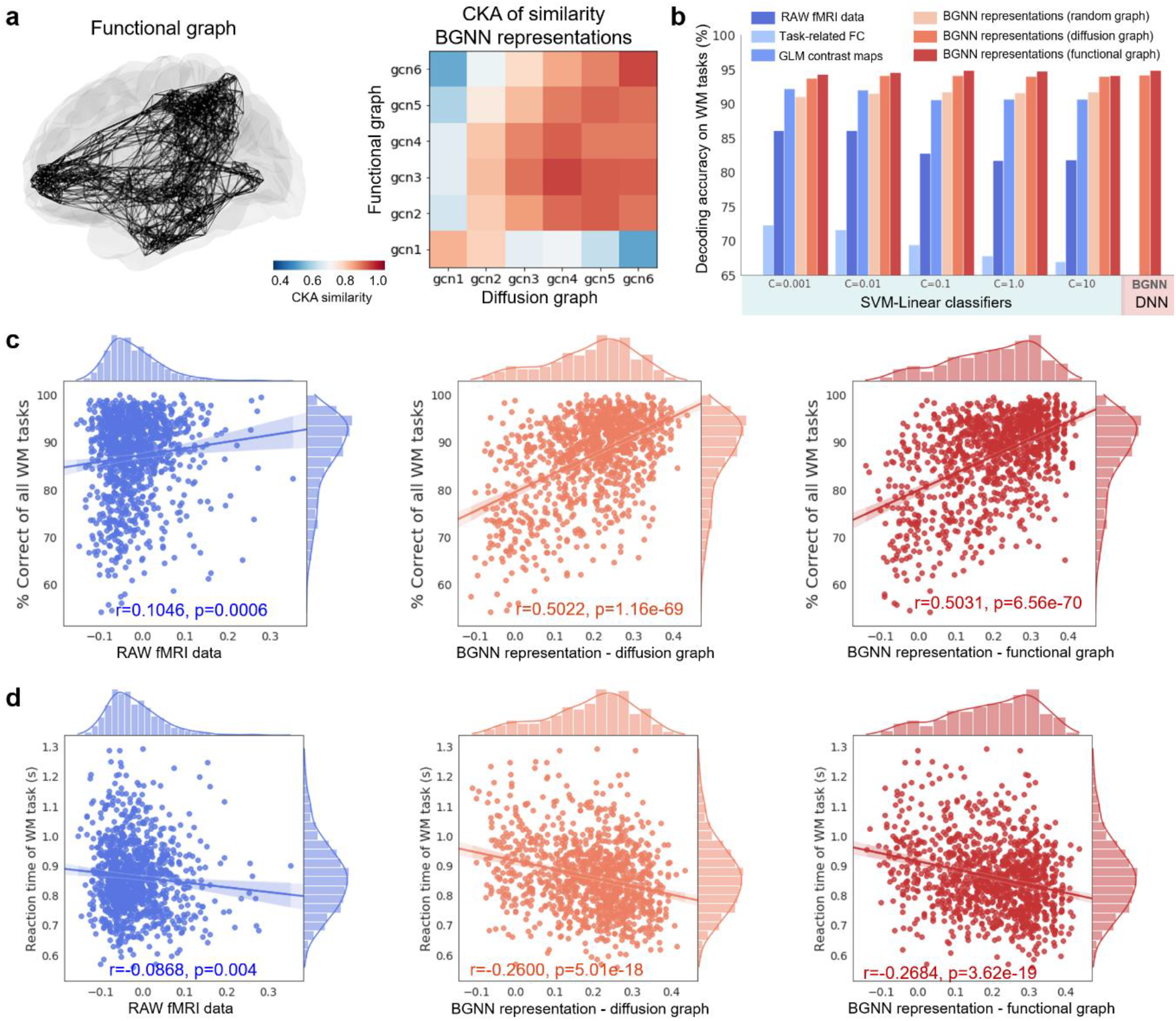
Interpretable representations of connectome-constrained BGNN improved the decoding of cognitive functions and the associations with human behaviors. **a)**, Similar high-level BGNN representations were captured by using empirical connectome priors derived from either resting-state functional connectivity (functional graph) or diffusion tractography (diffusion graph). **b)**, BGNN representations improved the decoding of WM tasks. Compared to the conventional GLM-derived contrast maps and raw fMRI data, BGNN representations showed much higher decoding accuracies regardless of the chosen classifiers, e.g. the linear classifiers like support vector machine classification (SVC) with different hyperparameters or deep learning models such as BGNN (followed by a two-layer feedforward network). Connectome-based BGNN representations (using functional or diffusion graphs) showed similar decoding performance and both outperformed the randomly connected graph. **c)** and **d)**, Connectome-based BGNN representations were strongly associated with participants’ in-scanner task performance, much better than the raw fMRI data (blue lines). Similar levels of behavioral associations for BGNN representations using functional (red lines) or diffusion connectome priors (orange lines).

Moreover, BGNN representations achieved higher decoding accuracies of cognitive tasks as compared to other neural representations, including raw fMRI data, task-related functional connectivity (Cai et al., 2014; Jiang et al., 2020) and GLM contrast maps (Fig.**5b**), regardless of choices for the linear and nonlinear classifier or its parameters. Interestingly, using human connectome priors derived from either functional or diffusion MRI (Fig. **5a**), the BGNN model learned similar middle-to-high-level abstract representations of cognitive processes. Similar decoding performance was achieved by using either connectome prior, both of which outperformed the randomly connected graph (Fig.**5b**).

#### Individual variation in BGNN representations associates with participants’ behavioral performance

Although mapping neural responses into a common representational space, BGNN representations still preserved the individual variability in cognitive processes by relating task-related neural representations of individual brains to participants’ in-scanner behavioral performance. Studies have shown that the task-specific effect or modularity of individual fMRI data was significantly associated with participants’ task performance in behaviors (Saggar et al., 2018). Here, by constructing the individual state-transition graph using BGNN representations rather than using raw fMRI data, we found much stronger associations between task-related neural representations and cognitive behaviors on a large healthy population (Fig.**5c** and **d**). Specifically, the segregation of memory load (*2bk-0bk*) was highly associated with individual behaviors in scanner (as shown in Fig.5-S1), including positive correlations with the average accuracy (Acc) on all WM tasks (*r* = 0.5031 *p* =6.56e-70), on 0back tasks (*r* = 0.4450, *p* =2.33e-53) and on 2back tasks (*r* =0.3966, *p* =9.67e-42), as well as negative correlations with the median reaction time (RT) on all WM tasks (*r* =-0.2684, *p* =3.62e-19), on 0back tasks (*r* =-0.3686, *p* =8,87e-36) and on 2back tasks (*r* =-0.1114, *p* =0.0001). Similar brain-behavioral associations were achieved by embedding BGNN representations using functional or diffusion connectome priors (Fig.**5c** and **d**). This analysis was done by using all subjects from the *HCP S1200* database (*N* =1074 of all subjects with available behavioral and imaging data for WM tasks). These significant correlations were sustained after controlling for the effect of confounds including age, gender, handedness and head motion (*r* =0.4659, *p* =5.74e-59 for Acc; *r* =-0.2552, *p* =2.0e-16 for RT).

Moreover, both the task-segregation effect of BGNN representations and their brain-behavioral associations were gradually strengthened as going deeper along the representational hierarchy of WM tasks (Fig.5-S2). Besides, the task-segregation effect of BGNN representations was significantly heritable in HCP twin populations (h^2^=0.3597, see Table S3 for all heritability estimates) and shared genetic influences with behavioral scores (*ρ*_*g*_ =0.80 and −0.39 respectively for Acc and RT, see Table1 for phenotypic and genetic correlations between BGNN representations and behavioral performance).

**Table 1.** Shared genetic influences in BGNN representations and behavioral scores for WM tasks. BGNN representations of WM tasks as well as the in-scanner behavioral performance were significantly heritable in HCP twin populations, after controlling for confounding effects of age, gender, handedness and head motion (as shown in Table S3). In order to quantify the shared genetic variance in brain-behavioral associations, we conducted bivariate genetic analyses between BGNN representations and behavioral performance, including the average accuracy (Acc) and reaction time (RT). Both genetic and phenotypic correlations reached a high-level of significance (FDR corrected). ***: *p* <0.001; **: *p* <0.01; *: p<0.05; .: p<0.1.

| | Phenotypic correlation<br>($\rho_p$) | Genetic correlation<br>($\rho_g$) |
| --- | --- | --- |
| WM_Task_Acc | 0.4659 *** | 0.7992 *** |
| WM_Task_2bk_Acc | 0.3716 *** | 0.7731 *** |
| WM_Task_0bk_Acc | 0.4189 *** | 0.8650 *** |
| WM_Task_RT | -0.2552 *** | -0.3895 ** |
| WM_Task_2bk_RT | -0.1173 *** | -0.2455 . |
| WM_Task_0bk_RT | -0.3408 *** | -0.4967 ** |

**Table 2.** Decoding high-order cognitive tasks at different timescales. We trained a series of single-domain decoders by using fMRI responses of each cognitive domain exclusively. Three circumstances in cognitive decoding were considered by using different lengths of time windows, including single-volume prediction (i.e. using TR=0.72s fMRI signals), using 10s fMRI signals (approximately the shortest duration among all task trials), as well as single-trial prediction. Note that, considering the delay effect of hemodynamic responses, in the single-volume prediction experiments, we only used fMRI volumes at least 6s after the task onset for model training and evaluation. In the single-trial prediction experiments, we used variable lengths of time windows in the decoding model, according to the maximum duration of a single task trial, for instance 12s for MOTOR tasks and 25s for WM tasks. Our results showed that longer time windows resulted in higher decoding accuracy, with the largest improvement found in the classification of WM tasks, i.e. F1-score increased from 0.76 to 0.94, followed by relational processing tasks, i.e. F1-score increasing from 0.79 to 0.90.

| Task Domains | #Subj | #Samples (number of single trials) | #Cond | Task dura. of a single trial (s) | Decoding accuracy (F1-score) |  |  |
| --- | --- | --- | --- | --- | --- | --- | --- |
|  |  |  |  |  | Single-volume prediction | 10s fMRI signals | Single-trial prediction |
| Working Memory | 1085 | 17,360 | 8 | 25 | 0.7646 | 0.8552 | 0.9414 |
| Relational Processing | 1043 | 12,516 | 2 | 16 | 0.7995 | 0.8550 | 0.9059 |
| Social Cognition | 1051 | 10,510 | 2 | 23 | 0.9186 | 0.9481 | 0.9644 |
| Language | 1051 | 16,816 | 2 | 12 | 0.9625 | 0.9825 | 0.9836 |
| Emotion | 1047 | 12,564 | 2 | 18 | 0.9760 | 0.9943 | 0.9944 |
| Motor | 1083 | 21,660 | 5 | 12 | 0.9267 | 0.9734 | 0.9801 |

### Reliable and biological meaningful salient features of BGNN

To understand the biological basis of BGNN, we conducted the saliency map analysis which demonstrated distinctive neural basis among cognitive tasks and captured robust representations across individual trials and subjects. The stability of saliency maps was evaluated by using repeated-measure ANOVA among 24 HCP subjects, controlling for the random effect of subjects and experimental trials. Only the salient brain regions that having high saliency values (>0.2) and showing a significant effect of task (*p*<0.001) were reported in the following analysis. Taking the Motor and WM tasks as examples, we detected highly consistent salient features across different trials and subjects (as shown in Fig.**6**). For the Motor tasks, we detected salient task-specific features in the sensorimotor cortex, e.g. area 5m (region label=36 in the Glasser’s atlas) selectively activated during foot movements, area 2 selectively activated during hand movements, area 6v selectively activated during tongue movements. Besides, we observed hemispheric symmetric patterns for the movements of left and right body parts (Fig.**6c**). For Working-Memory tasks, which involves both sensory perception and memory load, the decoding model learned salient features related to both aspects, i.e. distinction between 0back vs 2back tasks and the recognition of face vs place images (Fig.**2d**). Specifically, ParaHippocampal Area 1 (PHA1) and V4 Transitional Area (V4t) were selectively involved for the recognition of place images (repeated measure ANOVA, F-score=70.96 and 163.34, p-value=1.74e-8 and 6.21e-12 respectively for PHA1 and V4t), while Fusiform Face Complex (FFC) and Lateral Occipital Area 1 (LO1) were selectively engaged for the recognition of faces (F-score=57.75 and 91.47, p-value=1.02e-7 and 1.75e-9 respectively for FFC and LO1). On other hand, for both place and face images, Ventral Visual Complex (VVC) was more involved in 0back tasks than 2back tasks (F-score=39.86, p-value=2.0e-6) while Area 37 was selectively engaged in the 2back tasks (F-score=102.56, p-value=6.01e-10) when fixing the category of visual stimuli. Our results revealed that reliable representations were captured during cognitive decoding, which are not only biologically meaningful, e.g., engaging task-related brain regions, and more importantly show reliable and task-selective responses to cognitive tasks.

**Fig. 6.**
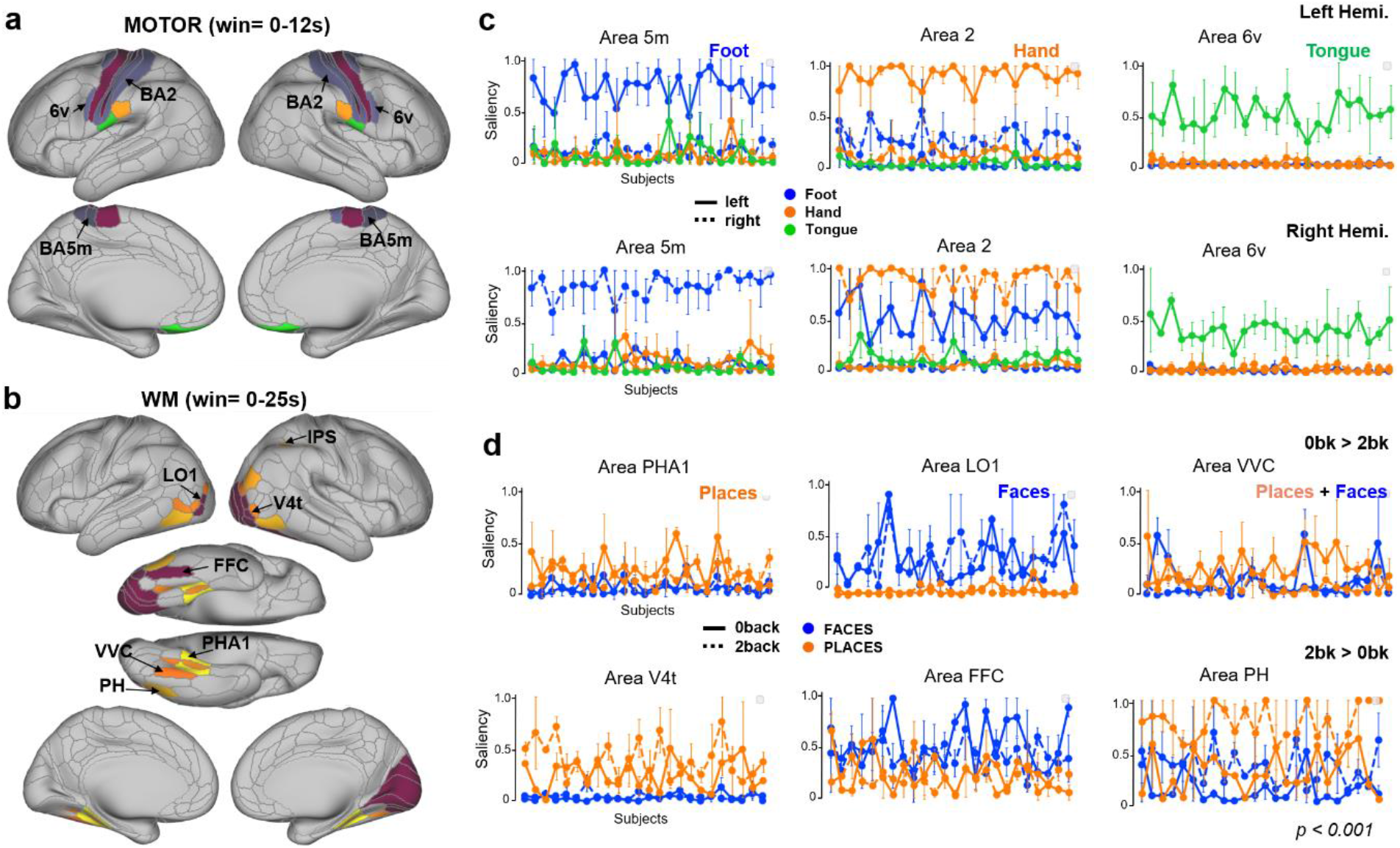
Salient BGNN features for the Motor and Working-memory tasks and their reliability. Only salient brain regions (saliency values>0.2, the full range of saliency is (0,1)) with a significant ‘task condition’ effect (p<0.001) was shown in **a**) and **b**) with the color scheme indicating different region id in Glasser’s atlas. We observed task-specific salient brain regions for Motor tasks (**c**), showing selective responses to the movement of foot (area 5m), hand (area 2) and tongue (area 6v), in solid lines for the movement of left side and in dashed lines for the right side. Symmetrical patterns of brain responses were detected in the salient regions in the both left (1^st^ row) and right hemisphere (2^nd^ row). We detected three sets of salient brain regions for WM tasks (**d**), showing selective responses to the image category, e.g. place (1^st^ column, in orange) and face images (2^nd^ column, in blue), or to memory load, e.g. 0back (solid lines) and 2back tasks (3^rd^ column, dashed lines). Error bars in the plots indicated the standard deviation of brain responses across repeated task trials within each subject.

## Discussion

In the present study, we proposed biologically-constrained graph neural networks (BGNNs) to model task-evoked brain dynamics by combining local and distributed brain activity through connectome-based constraints. By restricting the activity flow of cognitive tasks through anatomical or functional connections, BGNN revealed multilevel and multi-stage representations underpinning cognitive processes. At the low-level representation, BGNN uncovered a spatiotemporal decomposition of cognitive processes into multiple temporal stages and different patterns of spatial activation maps at each stage (e.g. *motor execution* and *motor preparation* for Motor tasks). At the high-level representation, BGNN learned inheritable and interpretable abstract representations of cognitive processes that improved inter-subject alignment in brain responses, enhanced cognitive decoding with high accuracy and fine timescales, and showed strong phenotypic and genetic correlations with individual behaviors (e.g. *correct responses* and *response time* of WM tasks). Moreover, the model uncovered a functional gradient in neural representations of WM, with a stepwise progression from sensory processing to cognitive control and towards behavioral abstraction, and revealed distinct neural substrates for the short-term memory of faces vs places, suggesting a privileged WM state of remembering faces. Together, these results demonstrate that, far from a black box, BGNNs lead to interpretable cognitive models and representational learning of human brain functions.

Our results revealed an important role of functional integration in cognitive processes, not only affecting the decoding of cognitive states but also changing the organizational principles of encoded brain representations. For segregated brain function like the motor processes, the modeling of within-network integration (K=1) is sufficient to achieve the optimal decoding performance and reveals a stable two-level hierarchy in neural representations (Fig. 6-S1c), namely the involvement of the sensorimotor cortex for motor execution and prefrontal regions for motor planning (Fig.**3**). The multilevel representations of Motor tasks coincided with previous findings showing a clear gradient of neural responses from preparation to execution in a sequential motor task (Orban et al., 2015) and prefrontal responses being predictive to body movements before execution (Ryun et al., 2014). For high-order cognition such as visual WM tasks, on the other hand, the modeling of between-network communication and functional integration (K>1) is critical to encode the multiscale, hierarchical representations of cognitive processes, namely image recognition, memory maintenance and representational abstraction (Fig.**2**). The three-level representations of WM were encoded in the responses of different sets of brain regions, consisting of the ventral visual stream, frontoparietal network regions, prefrontal and salience network regions, respectively (Fig.**2c**), following the cortical hierarchy transforming from sensory areas to the prefrontal cortex (Brincat et al., 2018). This finding of multilevel representations of WM tasks coincided with the literature on the gradual progression from low-level motor/sensory inputs to high-level abstract representations of WM along the posterior-to-frontal gradient (Christophel et al., 2017; Oh et al., 2019), indicating an important role of prefrontal cortex in the process of transforming sensory perception into behaviorally relevant representations (Brincat et al., 2018; Nee and D’Esposito, 2018; Oh et al., 2019).

The high-level abstract representations of WM tasks, captured by BGNNs with either anatomical or functional connectome priors, showed strong phenotypic and genetic correlations with individual behaviors, including both correct responses and reaction time of 0back and 2back WM tasks (Fig.5-S1). Interestingly, theses brain-behavior associations were gradually enhanced along representational hierarchy (Fig.5-S2), outperforming the predictive models of individual behaviors using either raw brain responses (Fig.5-S1) or resting-state functional connectivity (Yamashita et al., 2018). Our results suggest reliable behavioral abstraction and interpretable representational learning of WM by using connectome-constrained BGNN models.

Divergent brain mechanisms of the short-term memory were revealed for different types of visual stimuli, e.g., remembering faces vs places. Specifically, the retrieval of faces relies more on the heteromodal regions in the frontal and parietal cortices, while recognizing places mainly engages the unimodal regions in the ventral visual stream (Fig.**2c**). Consistently, participants also performed differently in behaviors among the two types of recognition tasks, i.e. showing higher accuracy and faster responses for the retrieval of faces than places (Fig.**2d** and Table S2). Our findings coincided with the theory of a privileged WM state of faces that showed improved accuracy and response time compared to non-faces (Brady et al., 2019; Lin et al., 2019). These findings suggest a differential cognitive state and distinct neural representations for the short-term memory of faces, possibly through the top-down modulation from prefrontal and parietal regions.

The present study focused on the interpretability and robustness of the GNN models, one of the main challenges for deep learning applications in neuroscience research (Thomas et al., 2021). In particular, we showed that connectome-constrained BGNNs extract biologically meaningful and task-specific salient features from brain responses (Figs. 6 and 7) and capture behaviorally relevant representations of cognitive functions showing strong phenotypic and genetic correlations with individual behavioral performance (Fig.**5** and Table 1). Firstly, the saliency map analysis confirmed the involvement of well-known task-related brain regions (Fig.**6-S2**), for instance, salient features in the sensorimotor cortex for motor execution (Penfield and Boldrey, 1937), the perisylvian language areas for language comprehension (Friederici, 2011) and the ventral visual stream for image recognition (Golarai et al., 2007). Most of these regions have been used as priors in previous MVPA studies, for instance, decoding faces vs objects by using brain activity in the ventral stream (Haxby et al., 2011). More importantly, the saliency map detected a broad set of brain areas that contribute to different temporal stages of cognitive processes (Fig.4 and Fig.4-S1). The temporal dynamics of cognitive processes but has been mostly ignored in previous fMRI studies, by either using meta-analytic approaches (Bartley et al., 2018; Rubin et al., 2017), or GLM-derived activation maps (Poldrack et al., 2009; Varoquaux et al., 2018). The recent work of Loula and colleagues (Loula et al., 2018) demonstrated the feasibility of decoding visual stimuli with short inter-stimuli intervals in fMRI acquisitions. A study from our group (Orban et al., 2015) revealed a gradient of task-evoked activations in a sequential motor task by decomposing brain responses into multiple stages of the motor process. In the current study, we observed a similar functional gradient in cognitive processes through a series of spatiotemporal decompositions of task-evoked brain responses, for instance, at the preparation and execution stage of a motor task (Figs. 3 and 4), and at the stages of cue, auditory processing and button pressing of a language task (Fig.4-S1). Specifically, the engagement of the sensorimotor cortex at the execution stage and the involvement of prefrontal regions at the preparation stage of Motor tasks has been reliably detected in our model (Fig. 4). The feasibility of such predictive model of movements using prefrontal signals before the execution stage has been demonstrated in previous studies, for instance, in both fMRI acquisitions in healthy participants (Orban et al., 2015) and electrocorticography (ECoG) recordings in epilepsy patients (Ryun et al., 2014). Our results suggest that brain regions showing high predictive power to cognitive functions and behaviors at the individual level may not follow the canonical HRF and thus may not be detected by conventional univariate analyses. Our study provides a better understanding of the neural dynamics underpinning cognitive processes and opens new opportunities to discover new brain mechanisms of cognitive functions in both spatial and temporal domains.

## Conclusion

In summary, we provide in-depth interpretations of connectome-constrained GNN decoding models and reveal the multilevel and multi-stage representations underpinning cognitive processes. At the low-level representation, BGNN uncovered a series of spatiotemporal decompositions of cognitive processes, including multiple processing stages in the temporal domain and different patterns of activation maps in the spatial domain. At the high-level representation, BGNN captured behaviorally relevant representations of cognitive functions that strongly associated with human behaviors at the individual level and were inheritable in a twin design. In particular, our findings uncovered a functional gradient in the neural representations of cognitive tasks, for instance, from motor planning to execution for Motor tasks, and a stepwise progression of WM from sensory processing to cognitive control and towards behavioral abstraction. The present work suggests the feasibility of an interpretable cognitive model by leveraging the inductive bias of human connectome priors in GNN models. With the in-depth interpretations and multilevel representations, the proposed framework may be applicable in many subfields of cognitive neuroscience, ranging from cognitive modeling to brain stimulation or even neuromodulation.

## Materials and Methods

### fMRI Datasets and Preprocessing

We used the block-design task-fMRI dataset from the Human Connectome Project S1200 release (https://db.humanconnectome.org/data/projects/HCP_1200). The minimal preprocessed fMRI data in CIFTI formats were selected. The preprocessing pipelines includes two steps (Glasser et al., 2013): 1) fMRIVolume pipeline generates “minimally preprocessed” 4D time-series (i.e. “.nii.gz” file) that includes gradient unwarping, motion correction, fieldmap-based EPI distortion correction, brain-boundary-based registration of EPI to structural T1-weighted scan, non-linear (FNIRT) registration into MNI152 space, and grand-mean intensity normalization. 2) fMRISurface pipeline projects fMRI data from the cortical gray matter ribbon onto the individual brain surface and then onto template surface meshes (i.e. “dtseries.nii” file), followed by surface-based smoothing using a geodesic Gaussian algorithm. Further details on fMRI data acquisition, task design and preprocessing can be found in (Barch et al., 2013; Glasser et al., 2013). The task fMRI database includes six cognitive domains, which are emotion, language, motor, relational, social, and working memory. In total, there are 21 different experimental conditions. The detailed description of the task paradigms as well as the selected cognitive domains can be found in (Barch et al., 2013; Zhang et al., 2021)

During Motor tasks, participants are presented with visual cues that ask them to either tap their fingers, or squeeze toes, or move the tongue. Each block of a movement type (hand, foot or tongue) is preceded by a 3s cue and lasts for 12s. In each of the two runs, there are 13 blocks in total, including 2 blocks of tongue movements, 4 of hand movements and 4 of foot movements, as well as 3 additional fixation blocks (15s) in the middle of each run.

The working-memory (WM) tasks involve two-levels of cognitive functions, with a combination of the category recognition task and N-Back memory task. Specifically, participants are presented with pictures of places, tools, faces and body parts. These 4 different stimulus types are presented in separate blocks, with half of the blocks using a 2back working memory task (recognizing the same image after two image presentations) and the other half using a 0back working memory task (recognizing a single image presented at the beginning of a block). Each of the two runs contains 8 task blocks and 4 fixation blocks (15s). Each task block consists of a 2.5s cue indicating the task type, followed by 10 task trials (2.5s each). For each task trial, the stimulus is presented for 2 seconds, followed by a 500 ms inter-task interval (ITI) when participants need to respond as target or not.

The language task consists of two conditions, i.e. story or mathematics, with variable duration of auditory stimuli. In the story trials, participants are instructed to passively listen to brief auditory stories (5-9 sentences) adapted from Aesop’s fables, followed by a two-alternative-choice question and response on the topic of the story. In the mathematical trials, participants are presented with a series of arithmetic operations, e.g. addition and subtraction, followed by a two-alternative-choice question and response about the result of the operations. Overall, the mathematical trials last around 12-15 seconds while the story trials lasts 25-30 seconds. In order to match the length of the two conditions, the mathematical trials are presented in pairs in the middle of the task, along with one additional trial at the end of the task.

### Connectome-constrained graph convolution on brain activity

A brain graph provides a network representation of the human brain by associating nodes with brain regions and defining edges via anatomical or functional connections (Bullmore and Sporns, 2009). We recently found that convolutional operations on the brain graph can be used to decode brain states among a large number of cognitive tasks (Zhang et al., 2021). Here, we proposed a more generalized form of graph convolution by using high-order Chebyshev polynomials and explored how different scales of functional integration affects the encoding and decoding of cognitive functions.

#### Step 1: Construction of brain graph

The decoding pipeline started with a weighted graph *𝒢* = (*𝒱, ε, 𝒲*), where *𝒱* is a parcellation of cerebral cortex into *N* regions, *ε* is a set of connections between each pair of brain regions, with its weights defined as *𝒲* = (w_*ij*_)_*i*=1. .*N,j*=1. .*N*_ Many alternative approaches can be used to build such brain graph *𝒢*, for instance using different brain parcellation schemes and constructing various types of brain connectomes (for a review, see (Bullmore and Sporns, 2009)). Here, we used Glasser’s multi-modal parcellation, consisting of 360 areas in the cerebral cortex, bounded by sharp changes in cortical architecture, function, connectivity, and topography (Glasser et al., 2016). The edges between each pair of nodes were estimated by calculating the group averaged resting-state functional connectivity (RSFC) based on minimal preprocessed resting-state fMRI data from *N* = 1080 HCP subjects (Glasser et al., 2013). Additional preprocessing steps were applied before the calculation of RSFC, including regressing out the signals from white matter and csf, and bandpass temporal filtering on frequencies between 0.01 to 0.1 HZ. Functional connectivity was calculated on individual brains using Pearson correlation and then normalized using Fisher z-transform before averaging among the entire group of subjects. The resulting functional graph characterizes the intrinsic functional organization of the human brain among HCP populations. An alternative graph was constructed from the whole-cortex probabilistic diffusion tractography based on HCP diffusion-weighted MRI data, with the edges indicating the average proportion of fiber tracts (streamlines) between the seed and target parcels (Rosen and Halgren, 2021). After that, a k-nearest-neighbor (k-NN) graph was built from both graphs by only connecting each node to its 8 neighbors with the highest connectivity strength.

#### Step 2: Mapping of task-evoked brain activity onto the graph

After the construction of the brain graph (i.e. defining brain parcels and edges), for each functional run and each subject, the preprocessed task-fMRI data was then mapped onto the set of brain parcels, resulting in a 2-dimensional time-series matrix. This time-series matrix was first split into multiple blocks of cognitive tasks according to fMRI paradigms and then cut into sets of time-series of the chosen window size (e.g. 10 second). Shorter time windows were discarded in the process. The remaining time-series were treated as independent data samples during model training. As a result, we generated a large number of fMRI time-series matrices from all cognitive domains, i.e. a short time-series with duration of *T* for each of *N*brain parcels *x* ∈ ℝ^*N*×*T*^. The entire dataset consists of over 1000 subjects for each cognitive domain (see Table S1 for detailed information), in total of 14,895 functional runs across the six cognitive domains, and 138,662 data samples of fMRI signals *x* ∈ ℝ^*N*×*T*^ when using a 10s time window (i.e. 15 functional volumes at TR=0.72s).

#### Step 3: Spatiotemporal graph convolutions using BGNN

Graph convolution relies on the graph Laplacian, which is a smooth operator characterizing the magnitude of signal changes between adjacent nodes. The normalized graph Laplacian is defined as:

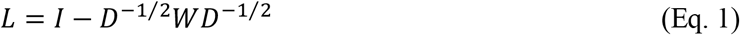

where *D* is a diagonal matrix of node degrees, *I* is the identity matrix, and *W* is the weight matrix. The eigendecomposition of Lapalcian matrix is defined as *L* = UΔU^*T*^, where U = (*u*_0_, *u*_1_, … *u*_*N*−1_) is the matrix of Laplacian eigenvectors and is also called graph Fourier modes, and Δ= diag(*λ*_0_, *λ*_1_, … *λ*_*N*−1_) is a diagonal matrix of the corresponding eigenvalues, specifying the frequency of the graph modes. In other words, the eigenvalues quantify the smoothness of signal changes on the graph, while the eigenvectors indicate the patterns of signal distribution on the graph.

For a signal *x* defined on graph, i.e. assigning a feature vector to each brain region, the convolution between the graph signal *x* ∈ ℝ^*N*×*T*^ and a graph filter *g*_*θ*_ ∈ ℝ^*N*×*T*^ based on graph *𝒢*, is defined as their element-wise Hadamard product in the spectral domain, i.e.:

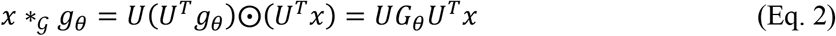

where *G*_*θ*_ = *diag*(*U*^*T*^*g*_*θ*_) and *θ* indicate a parametric model for graph convolution *g*_*θ*_, U = (*u*_0_, *u*_1_, … *u*_*N*−1_) is the matrix of Laplacian eigenvectors and *U*^*T*^*x* is projecting the graph signal onto the full spectrum of graph modes. To avoid calculating the spectral decomposition of the graph Laplacian, ChebNet convolution (Defferrard et al., 2016) uses a truncated expansion of the Chebychev polynomials, which are defined recursively by:

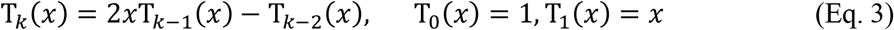

Consequently, the ChebNet graph convolution is defined as:

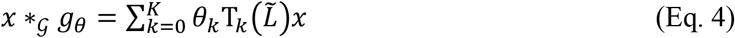

where 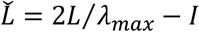 is a normalized version of graph Laplacian with *λ*_*max*_ being the largest eigenvalue, _*θk*_ is the model parameter to be learned at each order of the Chebychev polynomials. It has been proved that the ChebNet graph convolution was naturally *K*-localized in space by taking up to *K*th order Chebychev polynomials (Defferrard et al., 2016), which means that each ChebNet convolutional layer integrates the context of brain activity within a *K*-step neighborhood.

#### Step 4: The encoding-decoding model of brain responses

We proposed an encoding-decoding model based on ChebNet graph convolutions (Fig.**1**), consisting of 6 graph convolutional layers (6 BGNN layers) with 32 graph filters at each layer, followed by a flatten layer and 2 fully connected layers (256, 64 units). The encoding model takes in a short series of fMRI volumes as input, propagates brain activity within (K=1) and between (K>1) brain networks, and learns various shapes of temporal convolution kernels (*T* time points) as well as a rich set of spatial “brain activation” maps (*N* brain regions). The decoding model takes in the learned representations from the encoding model and predicts cognitive states via a 2-layer multilayer perceptron (MLP). The entire dataset was split into training (60%), validation (20%), test (20%) sets using a subject-specific split scheme, i.e. all fMRI data from the same subject being assigned to only one of the three sets. Approximately, the training set includes fMRI data from 700 unique subjects (depending on data availability for different cognitive tasks ranging from 1043 to 1085 subjects, see Table S1), with 176 subjects for validation set and 219 subjects for test set. The encoding-decoding model was jointly trained to predict the cognitive state from a short time window, e.g. 10s fMRI time-series. We used Adam as the optimizer with the initial learning rate as 0.0001 on all cognitive domains and saved the best model after 100 training epochs. Additional l2 regularization of 0.0005 on weights and a dropout rate of 0.5 was used to control model overfitting and the noise effect of fMRI signals. The implementation of the ChebNet graph convolution was based on PyTorch 1.1.0, and has been made publicly available in the repository: https://github.com/zhangyu2ustc/gcn_tutorial_test.git.

### Effects of K-order in ChebNet graph convolution

As stated in equation (4), the graph convolution can be rewritten as follows at different *K-*orders:

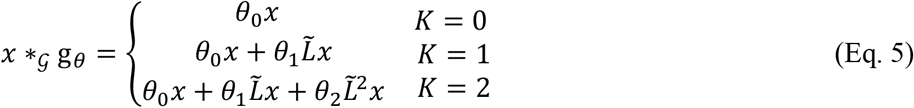

where 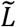 is a normalized version of graph Laplacian and _{*θ*_*k*_}*k*=1,2,. .*K*_ are model parameters to be trained. Specifically, *K=*0 indicates a global scaling factor on the input signal by treating each node independently, similar to the classical univariate analysis for brain activation detection; *K=*1 indicates information integration between the direct neighbors and the current node on the graph (i.e. integrating signals within the same network); *K=*2 indicates functional integration within a two-step neighborhood on the graph (i.e. integrating information from local area, within network and between networks). Thus, the choice of *K-*order controls the scale of the information integration on the graph. We explored different choices of *K*-order in ChebNet spanning over the list of [0,1,2,5,10] and found a significant boost in both brain decoding and representational learning by using high-order graph convolutions.

#### Similarity analysis of layer representations in BGNN

The BGNN model maps the spatiotemporal dynamics of fMRI brain activity onto a new representational space in the spectral domain. Different representations are learned at each BGNN layer by integrating activity flow within (K =1) and between networks (K >1). We analyzed the similarity of layer representations in BGNN by using centered kernel alignment (CKA) with a linear kernel. CKA was originally proposed to compare high-dimensional layer representations of deep neural networks, not only in the same network trained from different initializations, but also across different models (Kornblith et al., 2019). Here, we used CKA to evaluate the hierarchical organization of BGNN representations for both Motor and WM tasks. First, we extracted the learned representations from each layer using samples from the test set and reshaped the representations (*samples* × *brain regions* × *oime poinos*) into a 2D matrix *X* ∈ ℝ^*samples*×*oeaoures*^. Then, the linear CKA of two representation matrices *X* and *Y*, either from different layers or different models, was defined as:

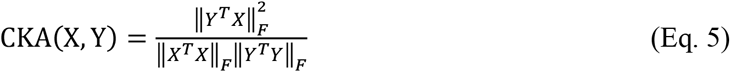

where 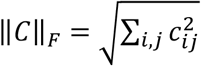 indicates the Frobenius norm of the cross-correlation matrix **C**. The CKA value was within the range [0,1], with its highest value at 1 (the same layer representation) and lowest at 0 (totally different layer representations). Next, a between-layer CKA matrix was calculated for each BGNN model and the hierarchical organization was revealed by using ward linkage.

### Projections of layer representations using t-SNE

For visualization purposes, we projected the high-dimensional layer representations (360*32 in our case) to a 2D space by using t-SNE (Maaten and Hinton, 2008). Based on the t-SNE projections, we calculated the modularity score among different task conditions as a measure of task segregation, representing the cost of brain state transition between tasks. It has been shown that the modularity score on the individual state-transition graph constructed from task-fMRI data was significantly associated with participants’ in-scanner task performance (Saggar et al., 2018). Here, we estimated the modularity score for both fMRI signals and layer representations of BGNN. Specifically, fMRI signals and layer representations were first mapped onto a 2D space by using t-SNE. Then, a k-NN graph (k=5) was constructed based on the coordinates of t-SNE projections by connecting each data sample with its five nearest neighbors in the 2D space. After that, the modularity score (*Q*) was calculated based on the partition of communities using task conditions (e.g. 0bk vs 2bk in WM tasks), with a high separation value indicating more edges (or similar representations) within the same task than expected by chance (Newman, 2006).

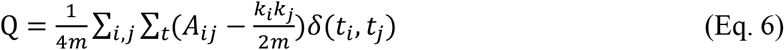

where *k*_*i*_ is the node degree of the kNN graph,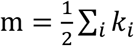 is the total number of edges, *o*_*ij*_ is the adjacent matrix, indicating whether node *i* and node *j* are connected in the kNN graph, and *δ*(*c*_*i*_, *c*_*j*_) indicates whether the two nodes belong to the same task. The task segregation index (*Q*) was within the range [-0.5,1], with the value close to 1 indicating a strong community structure in the BGNN representations of different task conditions. The task segregation was then correlated with participants’ in-scanner task performance, including averaged correct responses and reaction time during WM tasks.

### Saliency map analysis of the trained model

The saliency map analysis aims to locate which part of the brain contributes to the differentiation of cognitive tasks. We used a gradient approach named GuidedBackprop (Springenberg et al., 2014) to generate the saliency maps for each cognitive domain. Specifically, for the graph signal *X*^*l*^ of layer *l* and its gradient, *R*^*l*^ the overwritten gradient can be calculated as follows:

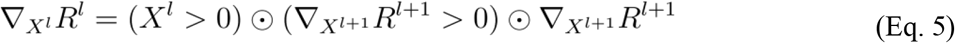

In order to generate the saliency map, we started from the output layer of a pre-trained model and used the above chain rule to propagate the gradients at each layer until reaching the input layer. This guided-backpropagation approach provides a high-resolution saliency for each data sample of fMRI signals *x* ∈ ℝ^*N*×*T*^. Then, a heatmap was calculated based on the saliency by taking the variance across all time steps for each parcel and normalizing it to the range [0,1], with its highest value at 1 (a dominant effect for task prediction) and lowest at 0 (no contribution to task prediction).

### Heritability analysis of brain representations

For the heritability estimates of brain responses of WM tasks, we used the Sequential Oligogenic Linkage Analysis Routines (SOLAR) Eclipse software package (http://www.nitrc.org/projects/se_linux). SOLAR relies on the maximum variance decomposition of the covariance matrix Ω for a pedigree:

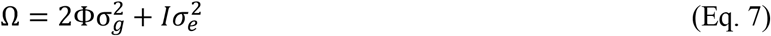

where 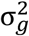 is the genetic variance due to the additive genetic factors, Φ is the kinship matrix representing the pairwise kinship coefficients among all individuals,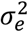 is the variance due to individual-specific environmental effects and measurement error, and *I* is an identity matrix. Narrow sense heritability is defined as the fraction of phenotypic variance 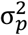 attributable to additive genetic factors: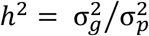. The significance of the heritability estimate is tested by comparing it to the model in which 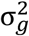 is constrained to zero. The heritability estimate was applied on 1074 subjects from HCP S1200 release with available behavioral and imaging data for WM tasks, which consist of 448 unique families, including 151 monozygotic-twin pairs, 92 dizygotic-twin pairs and 537 non-twin siblings. Prior to the heritability estimation, all phenotypes (brain and behavioral phenotypes) were adjusted for covariates including age, gender, handedness and head motion.

We further performed the bivariate genetic analyses to quantify the shared genetic variance and phenotypic correlation between brain responses and behavioral measures:

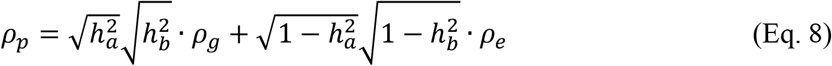

where *ρ*_*g*_ is the proportion of variability due to shared genetic effects and *ρ*_*e*_ is that due to the environment, while 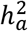 and 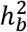 correspond to the narrow sense heritability for phenotypes *a* (representation of brain response) and *b* (behavioral scores).

## Acknowledgment

This work was partially supported by the Science and Technology Innovation 2030 - Brain Science and Brain-Inspired Intelligence Project (Grant No.2021ZD0200201, No.2022ZD0211500), Scientific Project of Zhejiang Lab (No.2022ND0AN01, No. 2022KI0AC02), Courtois foundation through the Courtois NeuroMod Project. PB is supported by a salary award of “Fonds de recherche du Québec - Santé”, chercheur boursier junior 2.

Data were provided by the Human Connectome Project, WU-Minn Consortium (Principal Investigators: David Van Essen and Kamil Ugurbil; 1U54MH091657) funded by the 16 NIH Institutes and Centers that support the NIH Blueprint for Neuroscience Research; and by the McDonnell Center for Systems Neuroscience at Washington University.

## Author contributions

Conceptualization: YZ, PB;

Methodology: YZ, PB;

Visualization: YZ, LF, PB;

Investigation: YZ, LF, TJ, AD, PB;

Writing—original draft: YZ, LF, TJ, AD, PB

Writing—review & editing: YZ, LF, TJ, AD, PB

## Ethics statement

Data were provided by the Human Connectome Project, WU-Minn Consortium (Principal Investigators: David Van Essen and Kamil Ugurbil; 1U54MH091657), with the approval of local ethics committee at Centre de recherche de l’Institut universitaire de gériatrie de Montréal (CRIUGM).

## Data and materials availability

We used publicly available dataset from the Human Connectome Project S1200 release, downloaded from https://db.humanconnectome.org/data/projects/HCP_1200. In total, fMRI data from 1095 unique subjects under six different task domains and resting-state were used in this study. The minimal preprocessed fMRI data of the CIFTI format were used, which maps individual fMRI time-series onto the standard surface template with 32k vertices per hemisphere. Our decoding pipeline, as well as the interpretations of BGNN models, were made publicly available in the following repository: https://github.com/zhangyu2ustc/gcn_tutorial_test.git

## References

Aguirre, G.K., Zarahn, E., D’Esposito, M., 1998. The Variability of Human, BOLD Hemodynamic Responses. NeuroImage 8, 360–369. https://doi.org/10.1006/nimg.1998.0369

Ariani, G., Pruszynski, J.A., Diedrichsen, J., 2022. Motor planning brings human primary somatosensory cortex into action-specific preparatory states. eLife 11, e69517. https://doi.org/10.7554/eLife.69517

Barch, D.M., Burgess, G.C., Harms, M.P., Petersen, S.E., Schlaggar, B.L., Corbetta, M., Glasser, M.F., Curtiss, S., Dixit, S., Feldt, C., Nolan, D., Bryant, E., Hartley, T., Footer, O., Bjork, J.M., Poldrack, R., Smith, S., Johansen-Berg, H., Snyder, A.Z., Van Essen, D.C., 2013. Function in the human connectome: Task-fMRI and individual differences in behavior. NeuroImage, Mapping the Connectome 80, 169–189. https://doi.org/10.1016/j.neuroimage.2013.05.033

Bartley, J.E., Boeving, E.R., Riedel, M.C., Bottenhorn, K.L., Salo, T., Eickhoff, S.B., Brewe, E., Sutherland, M.T., Laird, A.R., 2018. Meta-analytic evidence for a core problem solving network across multiple representational domains. Neurosci. Biobehav. Rev. 92, 318–337. https://doi.org/10.1016/j.neubiorev.2018.06.009

Bazeille, T., DuPre, E., Richard, H., Poline, J.-B., Thirion, B., 2021. An empirical evaluation of functional alignment using inter-subject decoding. NeuroImage 245, 118683. https://doi.org/10.1016/j.neuroimage.2021.118683

Brady, T.F., Alvarez, G.A., Störmer, V.S., 2019. The Role of Meaning in Visual Memory: Face-Selective Brain Activity Predicts Memory for Ambiguous Face Stimuli. J. Neurosci. 39, 1100–1108. https://doi.org/10.1523/JNEUROSCI.1693-18.2018

Brincat, S.L., Siegel, M., Nicolai, C. von, Miller, E.K., 2018. Gradual progression from sensory to task-related processing in cerebral cortex. Proc. Natl. Acad. Sci. 115, E7202–E7211. https://doi.org/10.1073/pnas.1717075115

Bullmore, E., Sporns, O., 2009. Complex brain networks: graph theoretical analysis of structural and functional systems. Nat. Rev. Neurosci. 10, 186–198. https://doi.org/10.1038/nrn2575

Cai, W., Ryali, S., Chen, T., Li, C.-S.R., Menon, V., 2014. Dissociable Roles of Right Inferior Frontal Cortex and Anterior Insula in Inhibitory Control: Evidence from Intrinsic and Task-Related Functional Parcellation, Connectivity, and Response Profile Analyses across Multiple Datasets. J. Neurosci. 34, 14652–14667. https://doi.org/10.1523/JNEUROSCI.3048-14.2014

Christophel, T.B., Hebart, M.N., Haynes, J.-D., 2012. Decoding the Contents of Visual Short-Term Memory from Human Visual and Parietal Cortex. J. Neurosci. 32, 12983– 12989. https://doi.org/10.1523/JNEUROSCI.0184-12.2012

Christophel, T.B., Klink, P.C., Spitzer, B., Roelfsema, P.R., Haynes, J.-D., 2017. The Distributed Nature of Working Memory. Trends Cogn. Sci. 21, 111–124. https://doi.org/10.1016/j.tics.2016.12.007

Defferrard, M., Bresson, X., Vandergheynst, P., 2016. Convolutional Neural Networks on Graphs with Fast Localized Spectral Filtering. Adv. Neural Inf. Process. Syst. 29.

D’Esposito, M., Postle, B.R., 2015. The cognitive neuroscience of working memory. Annu. Rev. Psychol. 66, 115–142. https://doi.org/10.1146/annurev-psych-010814-015031

Eriksson, D., Heiland, M., Schneider, A., Diester, I., 2021. Distinct dynamics of neuronal activity during concurrent motor planning and execution. Nat. Commun. 12, 5390. https://doi.org/10.1038/s41467-021-25558-8

Eriksson, J., Vogel, E.K., Lansner, A., Bergström, F., Nyberg, L., 2015. Neurocognitive Architecture of Working Memory. Neuron 88, 33–46. https://doi.org/10.1016/j.neuron.2015.09.020

Farroni, T., Johnson, M.H., Menon, E., Zulian, L., Faraguna, D., Csibra, G., 2005. Newborns’ preference for face-relevant stimuli: Effects of contrast polarity. Proc. Natl. Acad. Sci. 102, 17245–17250. https://doi.org/10.1073/pnas.0502205102

Friederici, A.D., 2011. The Brain Basis of Language Processing: From Structure to Function. Physiol. Rev. 91, 1357–1392. https://doi.org/10.1152/physrev.00006.2011

Gallivan, J.P., McLean, D.A., Valyear, K.F., Pettypiece, C.E., Culham, J.C., 2011. Decoding Action Intentions from Preparatory Brain Activity in Human Parieto-Frontal Networks. J. Neurosci. 31, 9599–9610. https://doi.org/10.1523/JNEUROSCI.0080-11.2011

Glasser, M.F., Coalson, T.S., Robinson, E.C., Hacker, C.D., Harwell, J., Yacoub, E., Ugurbil, K., Andersson, J., Beckmann, C.F., Jenkinson, M., Smith, S.M., Van Essen, D.C., 2016. A multi-modal parcellation of human cerebral cortex. Nature 536, 171–178. https://doi.org/10.1038/nature18933

Glasser, M.F., Sotiropoulos, S.N., Wilson, J.A., Coalson, T.S., Fischl, B., Andersson, J.L., Xu, J., Jbabdi, S., Webster, M., Polimeni, J.R., Van Essen, D.C., Jenkinson, M., 2013. The minimal preprocessing pipelines for the Human Connectome Project. NeuroImage, Mapping the Connectome 80, 105–124. https://doi.org/10.1016/j.neuroimage.2013.04.127

Golarai, G., Ghahremani, D.G., Whitfield-Gabrieli, S., Reiss, A., Eberhardt, J.L., Gabrieli, J.D., Grill-Spector, K., 2007. Differential development of high-level visual cortex correlates with category-specific recognition memory. Nat. Neurosci. 10, 512–522. https://doi.org/10.1038/nn1865

Groen, I.I., Greene, M.R., Baldassano, C., Fei-Fei, L., Beck, D.M., Baker, C.I., 2018. Distinct contributions of functional and deep neural network features to representational similarity of scenes in human brain and behavior. eLife 7, e32962. https://doi.org/10.7554/eLife.32962

Guntupalli, J.S., Feilong, M., Haxby, J.V., 2018. A computational model of shared fine-scale structure in the human connectome. PLOS Comput. Biol. 14, e1006120. https://doi.org/10.1371/journal.pcbi.1006120

Guntupalli, J.S., Hanke, M., Halchenko, Y.O., Connolly, A.C., Ramadge, P.J., Haxby, J.V., 2016. A Model of Representational Spaces in Human Cortex. Cereb. Cortex N. Y. N 1991 26, 2919–2934. https://doi.org/10.1093/cercor/bhw068

Harrison, S.A., Tong, F., 2009. Decoding reveals the contents of visual working memory in early visual areas. Nature 458, 632–635. https://doi.org/10.1038/nature07832

Haxby, J.V., 2012. Multivariate pattern analysis of fMRI: The early beginnings. NeuroImage 62, 852–855. https://doi.org/10.1016/j.neuroimage.2012.03.016

Haxby, J.V., Connolly, A.C., Guntupalli, J.S., 2014. Decoding Neural Representational Spaces Using Multivariate Pattern Analysis. Annu. Rev. Neurosci. 37, 435–456. https://doi.org/10.1146/annurev-neuro-062012-170325

Haxby, J.V., Guntupalli, J.S., Connolly, A.C., Halchenko, Y.O., Conroy, B.R., Gobbini, M.I., Hanke, M., Ramadge, P.J., 2011. A Common, High-Dimensional Model of the Representational Space in Human Ventral Temporal Cortex. Neuron 72, 404–416. https://doi.org/10.1016/j.neuron.2011.08.026

Haxby, J.V., Guntupalli, J.S., Nastase, S.A., Feilong, M., 2020. Hyperalignment: Modeling shared information encoded in idiosyncratic cortical topographies. eLife 9, e56601. https://doi.org/10.7554/eLife.56601

Hou, Y., Jia, S., Zhang, S., Lun, X., Shi, Y., Li, Y., Yang, H., Zeng, R., Lv, J., 2020. Deep Feature Mining via Attention-based BiLSTM-GCN for Human Motor Imagery Recognition. ArXiv200500777 Cs Eess.

Huth, A.G., Nishimoto, S., Vu, A.T., Gallant, J.L., 2012. A continuous semantic space describes the representation of thousands of object and action categories across the human brain. Neuron 76, 1210–1224. https://doi.org/10.1016/j.neuron.2012.10.014

Jiang, R., Zuo, N., Ford, J.M., Qi, S., Zhi, D., Zhuo, C., Xu, Y., Fu, Z., Bustillo, J., Turner, J.A., Calhoun, V.D., Sui, J., 2020. Task-induced brain connectivity promotes the detection of individual differences in brain-behavior relationships. NeuroImage 207, 116370. https://doi.org/10.1016/j.neuroimage.2019.116370

Kell, A.J.E., Yamins, D.L.K., Shook, E.N., Norman-Haignere, S.V., McDermott, J.H., 2018. A Task-Optimized Neural Network Replicates Human Auditory Behavior, Predicts Brain Responses, and Reveals a Cortical Processing Hierarchy. Neuron 98, 630-644.e16. https://doi.org/10.1016/j.neuron.2018.03.044

Kornblith, S., Norouzi, M., Lee, H., Hinton, G., 2019. Similarity of Neural Network Representations Revisited. ArXiv190500414 Cs Q-Bio Stat.

Kriegeskorte, N., Douglas, P.K., 2018. Cognitive computational neuroscience. Nat. Neurosci. 21, 1148–1160. https://doi.org/10.1038/s41593-018-0210-5

Levakov, G., Faskowitz, J., Avidan, G., Sporns, O., 2021. Mapping individual differences across brain network structure to function and behavior with connectome embedding. NeuroImage 242, 118469. https://doi.org/10.1016/j.neuroimage.2021.118469

Li, X., Zhou, Y., Dvornek, N., Zhang, M., Gao, S., Zhuang, J., Scheinost, D., Staib, L.H., Ventola, P., Duncan, J.S., 2021. BrainGNN: Interpretable Brain Graph Neural Network for fMRI Analysis. Med. Image Anal. 74, 102233. https://doi.org/10.1016/j.media.2021.102233

Lin, H., Li, W., Carlson, S., 2019. A Privileged Working Memory State and Potential Top-Down Modulation for Faces, Not Scenes. Front. Hum. Neurosci. 13, 2. https://doi.org/10.3389/fnhum.2019.00002

Lin, Y., Yang, D., Hou, J., Yan, C., Kim, M., Laurienti, P.J., Wu, G., 2021. Learning dynamic graph embeddings for accurate detection of cognitive state changes in functional brain networks. NeuroImage 230, 117791. https://doi.org/10.1016/j.neuroimage.2021.117791

Llera, A., Wolfers, T., Mulders, P., Beckmann, C.F., 2019. Inter-individual differences in human brain structure and morphology link to variation in demographics and behavior [WWW Document]. eLife. https://doi.org/10.7554/eLife.44443

Loula, J., Varoquaux, G., Thirion, B., 2018. Decoding fMRI activity in the time domain improves classification performance. NeuroImage, New advances in encoding and decoding of brain signals 180, 203–210. https://doi.org/10.1016/j.neuroimage.2017.08.018

Maaten, L. van der, Hinton, G., 2008. Visualizing Data using t-SNE. J. Mach. Learn. Res. 9, 2579–2605.

Messinger, A., Cirillo, R., Wise, S.P., Genovesio, A., 2021. Separable neuronal contributions to covertly attended locations and movement goals in macaque frontal cortex. Sci. Adv. 7, eabe0716. https://doi.org/10.1126/sciadv.abe0716

Mesulam, M.M., 1998. From sensation to cognition. Brain 121, 1013–1052. https://doi.org/10.1093/brain/121.6.1013

Mitchell, T.M., Shinkareva, S.V., Carlson, A., Chang, K.-M., Malave, V.L., Mason, R.A., Just, M.A., 2008. Predicting Human Brain Activity Associated with the Meanings of Nouns. Science 320, 1191–1195. https://doi.org/10.1126/science.1152876

Nakai, T., Nishimoto, S., 2020. Quantitative models reveal the organization of diverse cognitive functions in the brain. Nat. Commun. 11, 1142. https://doi.org/10.1038/s41467-020-14913-w

Naselaris, T., Olman, C.A., Stansbury, D.E., Ugurbil, K., Gallant, J.L., 2015. A voxel-wise encoding model for early visual areas decodes mental images of remembered scenes. NeuroImage 105, 215–228. https://doi.org/10.1016/j.neuroimage.2014.10.018

Nee, D.E., D’Esposito, M., 2018. The Representational Basis of Working Memory, in: Clark, R.E., Martin, S.J. (Eds.), Behavioral Neuroscience of Learning and Memory, Current Topics in Behavioral Neurosciences. Springer International Publishing, Cham, pp. 213–230. https://doi.org/10.1007/7854_2016_456

Neumann, J., Lohmann, G., Zysset, S., von Cramon, D.Y., 2003. Within-subject variability of BOLD response dynamics. NeuroImage 19, 784–796. https://doi.org/10.1016/S1053-8119(03)00177-0

Newman, M.E.J., 2006. Modularity and community structure in networks. Proc. Natl. Acad. Sci. 103, 8577–8582. https://doi.org/10.1073/pnas.0601602103

Nguyen, A., Yosinski, J., Clune, J., 2019. Understanding Neural Networks via Feature Visualization: A survey (No. arXiv:1904.08939). arXiv. https://doi.org/10.48550/arXiv.1904.08939

Nishida, S., Nishimoto, S., 2018. Decoding naturalistic experiences from human brain activity via distributed representations of words. NeuroImage, New advances in encoding and decoding of brain signals 180, 232–242. https://doi.org/10.1016/j.neuroimage.2017.08.017

Nishimoto, S., Vu, A.T., Naselaris, T., Benjamini, Y., Yu, B., Gallant, J.L., 2011. Reconstructing visual experiences from brain activity evoked by natural movies. Curr. Biol. 21, 1641–1646. https://doi.org/10.1016/j.cub.2011.08.031

Norman-Haignere, S., Kanwisher, N.G., McDermott, J.H., 2015. Distinct Cortical Pathways for Music and Speech Revealed by Hypothesis-Free Voxel Decomposition. Neuron 88, 1281–1296. https://doi.org/10.1016/j.neuron.2015.11.035

Oh, B.-I., Kim, Y.-J., Kang, M.-S., 2019. Ensemble representations reveal distinct neural coding of visual working memory. Nat. Commun. 10, 5665. https://doi.org/10.1038/s41467-019-13592-6

Orban, P., Doyon, J., Petrides, M., Mennes, M., Hoge, R., Bellec, P., 2015. The Richness of Task-Evoked Hemodynamic Responses Defines a Pseudohierarchy of Functionally Meaningful Brain Networks. Cereb. Cortex N. Y. N 1991 25, 2658–2669. https://doi.org/10.1093/cercor/bhu064

Penfield, W., Boldrey, E., 1937. SOMATIC MOTOR AND SENSORY REPRESENTATION IN THE CEREBRAL CORTEX OF MAN AS STUDIED BY ELECTRICAL STIMULATION1. Brain 60, 389–443. https://doi.org/10.1093/brain/60.4.389

Pinal, D., Zurrón, M., Díaz, F., 2014. Effects of load and maintenance duration on the time course of information encoding and retrieval in working memory: from perceptual analysis to post-categorization processes. Front. Hum. Neurosci. 8, 165. https://doi.org/10.3389/fnhum.2014.00165

Poldrack, R.A., Halchenko, Y., Hanson, S.J., 2009. Decoding the Large-Scale Structure of Brain Function by Classifying Mental States Across Individuals. Psychol. Sci. 20, 1364–1372. https://doi.org/10.1111/j.1467-9280.2009.02460.x

Pulvermüller, F., Tomasello, R., Henningsen-Schomers, M.R., Wennekers, T., 2021. Biological constraints on neural network models of cognitive function. Nat. Rev. Neurosci. 22, 488–502. https://doi.org/10.1038/s41583-021-00473-5

Riggall, A.C., Postle, B.R., 2012. The Relationship between Working Memory Storage and Elevated Activity as Measured with Functional Magnetic Resonance Imaging. J. Neurosci. 32, 12990–12998. https://doi.org/10.1523/JNEUROSCI.1892-12.2012

Rosen, B.Q., Halgren, E., 2021. A Whole-Cortex Probabilistic Diffusion Tractography Connectome. eNeuro 8. https://doi.org/10.1523/ENEURO.0416-20.2020

Rubin, T.N., Koyejo, O., Gorgolewski, K.J., Jones, M.N., Poldrack, R.A., Yarkoni, T., 2017. Decoding brain activity using a large-scale probabilistic functional-anatomical atlas of human cognition. PLOS Comput. Biol. 13, e1005649. https://doi.org/10.1371/journal.pcbi.1005649

Ryun, S., Kim, J.S., Lee, S.H., Jeong, S., Kim, S.-P., Chung, C.K., 2014. Movement Type Prediction before Its Onset Using Signals from Prefrontal Area: An Electrocorticography Study. BioMed Res. Int. 2014, e783203. https://doi.org/10.1155/2014/783203

Saggar, M., Sporns, O., Gonzalez-Castillo, J., Bandettini, P.A., Carlsson, G., Glover, G., Reiss, A.L., 2018. Towards a new approach to reveal dynamical organization of the brain using topological data analysis. Nat. Commun. 9, 1399. https://doi.org/10.1038/s41467-018-03664-4

Sato, W., Yoshikawa, S., 2013. Recognition memory for faces and scenes. J. Gen. Psychol. 140, 1–15. https://doi.org/10.1080/00221309.2012.710275

Shi, X., Lv, F., Seng, D., Zhang, J., Chen, J., Xing, B., 2020. Visualizing and understanding graph convolutional network. Multimed. Tools Appl. https://doi.org/10.1007/s11042-020-09885-4

Sligte, I.G., van Moorselaar, D., Vandenbroucke, A.R.E., 2013. Decoding the Contents of Visual Working Memory: Evidence for Process-Based and Content-Based Working Memory Areas? J. Neurosci. 33, 1293–1294. https://doi.org/10.1523/JNEUROSCI.4860-12.2013

Springenberg, J.T., Dosovitskiy, A., Brox, T., Riedmiller, M., 2014. Striving for Simplicity: The All Convolutional Net.

Stansbury, D.E., Naselaris, T., Gallant, J.L., 2013. Natural Scene Statistics Account for the Representation of Scene Categories in Human Visual Cortex. Neuron 79, 1025– 1034. https://doi.org/10.1016/j.neuron.2013.06.034

Tang, H., Qi, X.-L., Riley, M.R., Constantinidis, C., 2019. Working memory capacity is enhanced by distributed prefrontal activation and invariant temporal dynamics. Proc. Natl. Acad. Sci. 116, 7095–7100. https://doi.org/10.1073/pnas.1817278116

Thomas, A.W., Ré, C., Poldrack, R.A., 2021. Challenges for cognitive decoding using deep learning methods. ArXiv210806896 Cs Stat.

Van Essen, D.C., Smith, S.M., Barch, D.M., Behrens, T.E.J., Yacoub, E., Ugurbil, K., 2013. The WU-Minn Human Connectome Project: An overview. NeuroImage, Mapping the Connectome 80, 62–79. https://doi.org/10.1016/j.neuroimage.2013.05.041

Varoquaux, G., Schwartz, Y., Poldrack, R.A., Gauthier, B., Bzdok, D., Poline, J.-B., Thirion, B., 2018. Atlases of cognition with large-scale human brain mapping. PLOS Comput. Biol. 14, e1006565. https://doi.org/10.1371/journal.pcbi.1006565

Wang, D., Buckner, R.L., Fox, M.D., Holt, D.J., Holmes, A.J., Stoecklein, S., Langs, G., Pan, R., Qian, T., Li, K., Baker, J.T., Stufflebeam, S.M., Wang, K., Wang, X., Hong, B., Liu, H., 2015. Parcellating cortical functional networks in individuals. Nat. Neurosci. 18, 1853–1860. https://doi.org/10.1038/nn.4164

Xu, Y., Vaziri-Pashkam, M., 2021. Limits to visual representational correspondence between convolutional neural networks and the human brain. Nat. Commun. 12, 2065. https://doi.org/10.1038/s41467-021-22244-7

Yamashita, M., Yoshihara, Y., Hashimoto, R., Yahata, N., Ichikawa, N., Sakai, Y., Yamada, T., Matsukawa, N., Okada, G., Tanaka, S.C., Kasai, K., Kato, N., Okamoto, Y., Seymour, B., Takahashi, H., Kawato, M., Imamizu, H., 2018. A prediction model of working memory across health and psychiatric disease using whole-brain functional connectivity. eLife 7, e38844. https://doi.org/10.7554/eLife.38844

Zhang, Y., Bellec, P., 2020. Transferability of Brain decoding using Graph Convolutional Networks. bioRxiv 2020.06.21.163964. https://doi.org/10.1101/2020.06.21.163964

Zhang, Y., Farrugia, N., Bellec, P., 2022. Deep learning models of cognitive processes constrained by human brain connectomes. Med. Image Anal. 102507. https://doi.org/10.1016/j.media.2022.102507

Zhang, Y., Tetrel, L., Thirion, B., Bellec, P., 2021. Functional annotation of human cognitive states using deep graph convolution. NeuroImage 231, 117847. https://doi.org/10.1016/j.neuroimage.2021.117847

