## supplementary figures for "Interpretable brain decoding from sensations to cognition to action: graph neural networks reveal the representational hierarchy of human cognition"

**Running Title: Interpretable cognitive modeling using GNN**

**Authors:** Yu Zhang*, Lingzhong Fan, Tianzi Jiang, Alain Dagher and Pierre Bellec*

*** Corresponding Author:**

**Yu Zhang**, Research Institute of Artificial Intelligence, Zhejiang Lab.

**Pierre Bellec**, Department of Psychology, University de Montreal.

**This PDF file includes:**

Eleven Supplementary Figures: **Fig. 2-S1** to **Fig. 6-S2**

Three Supplementary Tables: **Table S1** to **Table S3**

### Supplementary results

#### Decoding multidomain cognitive functions

The proposed model was also applied to other cognitive domains acquired from HCP task-fMRI datasets (data description in Table S1). Using 10s of fMRI recordings, the model successfully identified the six cognitive domains with a test accuracy of 96%. The model achieved high decoding accuracies at multiple temporal resolutions, ranging from prediction on a single volume towards the entire task block/trial (see Table 2). In order to validate that the decoding model used biological meaningful features, we generated the saliency maps by propagating the non-negative gradients backwards to the input layer (Springenberg et al., 2014). We detected different sets of salient brain regions for each cognitive domain (as shown in Fig.**6-S2**), for instance the involvement of the somatosensory cortex for motor execution (MOTOR), the engagement of perisylvian language areas for language comprehension (LANGUAGE), and the association of the ventral visual stream for visual processing and pattern matching (WM, RELATIONAL and EMOTION). Moreover, we found that the scale of functional integration (specified by the *K*-order in ChebNet) showed a big impact on the decoding performance (Fig. 2-S3b), with different sensitivity levels across cognitive domains (Fig. 2-S3a). For instance, for the Motor task, the decoding model did not gain from between-network communication when using a high-order model and captured a stable two-level hierarchy in the BGNN representations at different Ks (Fig. 6-S1c). By contrast, the decoding of WM tasks gradually improved as increasing K and reaching the plateau after $K > 5$. Coincidingly, the three-level hierarchy of WM tasks was only captured in the high-order models but was broken in the ChebNet-*K*1 model (Fig. 6-S1d). Specifically, compared to the ChebNet-*K*5 model, the

ChebNet-*K*1 model successfully captured the low-level features in early BGNN layers, but failed to capture the double-dissociation between memory load and image category, e.g., differential representational states for remembering faces and places, in high-level representations (Fig.2-S1 and Fig.2-S4). Our findings coincided with the notion of functional segregation and integration in brain cognition (Bressler and Menon, 2010), for instance, within-network communication is essential for motor execution, whereas integrative, between-network communication is critical for visual working memory (Cohen and D'Esposito, 2016).

#### Variable sensitivity to the K-order uncovers different organizational principles in cognitive processes

The choice of *K*-order in ChebNet impacts the scale of information integration, by taking into account multilevel integration of neural dynamics at each graph convolutional layer, ranging from localized brain areas (K=0) to spatially distributed regions within the same network (K=1) and towards inter-connected brain networks (K>1). The choice of *K*-order not only showed an impact on the decoding performance, but also changed the hierarchy of feature representations learned in each BGNN layer.

First of all, the decoding of six cognitive domains significantly impacted by the choice of *K*-order in ChebNet, indicating a faster convergence speed as well as higher decoding accuracy when using high-order models (Fig. 6-S1b). Significant improvements in decoding were detected between K=1 (integration of brain activity within the same network) and K>1 (between-network communication) (test accuracy = 93% vs 96% respectively for K=1 and K>1), significantly boosted compared to the localized decoding model (test accuracy = 83% for K=0). Second, variable sensitivity to the *K*-order was detected among different cognitive domains (Fig. 6-S1a). Specifically, for the Motor task, the decoding performance showed no improvement when increasing *K*, which means no gain from between-network communication during motor execution. Coinciding with this, the hierarchical organization of layer representations in the Motor task showed a very stable bipartition pattern when increasing K, i.e. low- and high-level features (as shown in Fig. 6-S1b). By contrast, the decoding of WM tasks gradually improved as increasing K and reaching the plateau after $K > 5$, which means that between-network communication and high-order integration plays an important role in WM, especially for distinguishing between 0back and 2back tasks. Interestingly, the hierarchical organizational structure in WM (as shown in Fig. 6-S1c) started with three isolated clusters at K=1, gradually fused the representations by filling the gaps between neighboring layers, and converged to a stable tripartite organization at K=5 (i.e. low-, middle- and high-level representations). Further increase in the K-order did not change this organization but instead expanded the middle-level through encoding redundant hidden representations. Our results indicated that the variable sensitivity to the choice of *K*-order may uncover distinct organizational principles in cognitive processes, for instance, localized information processing within the motor and sensory cortex for motor execution, while complex forms of functional interaction and information integration across multiple brain systems/networks during WM tasks.

#### Functional integration in Working-memory tasks and segregation in Motor tasks

In order to evaluate the importance of functional segregation and integration for cognitive decoding, we conducted a systematic analysis on the decoding models at different *K*-orders and calculated the similarity of learned representations between BGNN models. We used the ChebNet-$K$5 model as the reference model for the similarity analysis.

For Motor tasks, the ChebNet-$K$1 model already captured the low-to-high-level organization in BGNN representations (Fig. 6-S1c). Further increase in K did not change this organization but only caused redundant representations in the high-level features (average similarity with gcn6 in gcn3-gcn5 is CKA=0.78 and 0.92 for ChebNet-$K$1 and ChebNet-$K$5). A direct comparison between the two models (3rd row and 1st column in Fig. 6-S2b) revealed that, compared to the ChebNet-$K$5 model, the ChebNet-$K$1 model captured highly similar low-level features (CKA=0.92 for gcn1 when comparing between ChebNet-$K$1 and ChebNet-$K$5) and learned closely related high-level representations (CKA=0.84 for gcn6 between the two models). However, very different hidden representations were learned in the middle layers between the two models (averaged CKA=0.70 for gcn2 to gcn5). These results indicated that the highly segregated brain function, such as the sensory and motor tasks, did not involve high-level of information integration, but rather relied on neural transmission of brain activity within a local area or segregated networks.

On the other hand, for the Working Memory tasks, the ChebNet-$K$5 model captured a nice disassociation between low-level features (gcn1 to gcn2), hidden representations (gcn3 to gcn4), and high-level features (gcn5 to gcn6). Such hierarchical organization was broken in the ChebNet-$K$1 model due to poor between-layer communication (i.e. big gaps in the representations between neighboring layers, CKA=0.98 and 0.69 for within- and between-level similarity in ChebNet-$K$1). Moreover, the ChebNet-$K$1 model successfully captured the low-level features by showing high similarity to ChebNet-$K$5 in the first two ChebNet layers, but it was not capable of encoding high-level representations in the last ChebNet layer (3rd row and 1st column in Fig. 6-S2c, compared between ChebNet-$K$1 and ChebNet-$K$5, CKA=0.93 for gcn1, 0.88 for gcn2, 0.74 for gcn6). By contrast, the ChebNet-$K$10 model learned very similar representations in the low, middle and high ChebNet layers as in ChebNet-K5 (4th row and 3rd column in Fig. 6-S2b, compared between ChebNet-$K$5 and ChebNet-$K$10, CKA=0.94 for gcn1, 0.90 for gcn6, average CKA=0.90 for gcn3 to gcn5). These results indicated that the high-order cognitive functions required a large scale of information propagation and integration on the brain graph, not only involving the local connections within a specific brain network ($K = 1$) but also engaging the long-range connections across multiple networks ($K \geq 5$).

#### Heritability of BGNN representations of brain responses

The segregation of BGNN representations for WM tasks was significantly heritable in HCP twin population ($h^2$ =0.3597 and 0.3493 for BGNN representations learned on the functional and diffusion graphs). Higher heritability levels were detected for in-scanner behavioral performance during WM tasks ($h^2$ =0.5624 and 0.4118 for average accuracy and reaction time in WM tasks, see Table S3 for all heritability estimates). Moreover, we found significantly shared genetic variance in BGNN representations and behavioral scores ($\rho_g$ =0.80 and -0.39 respectively for the average accuracy and reaction time, see Table 1 for shared genetic influences in brain-behavioral associations).

### Supplementary Figures

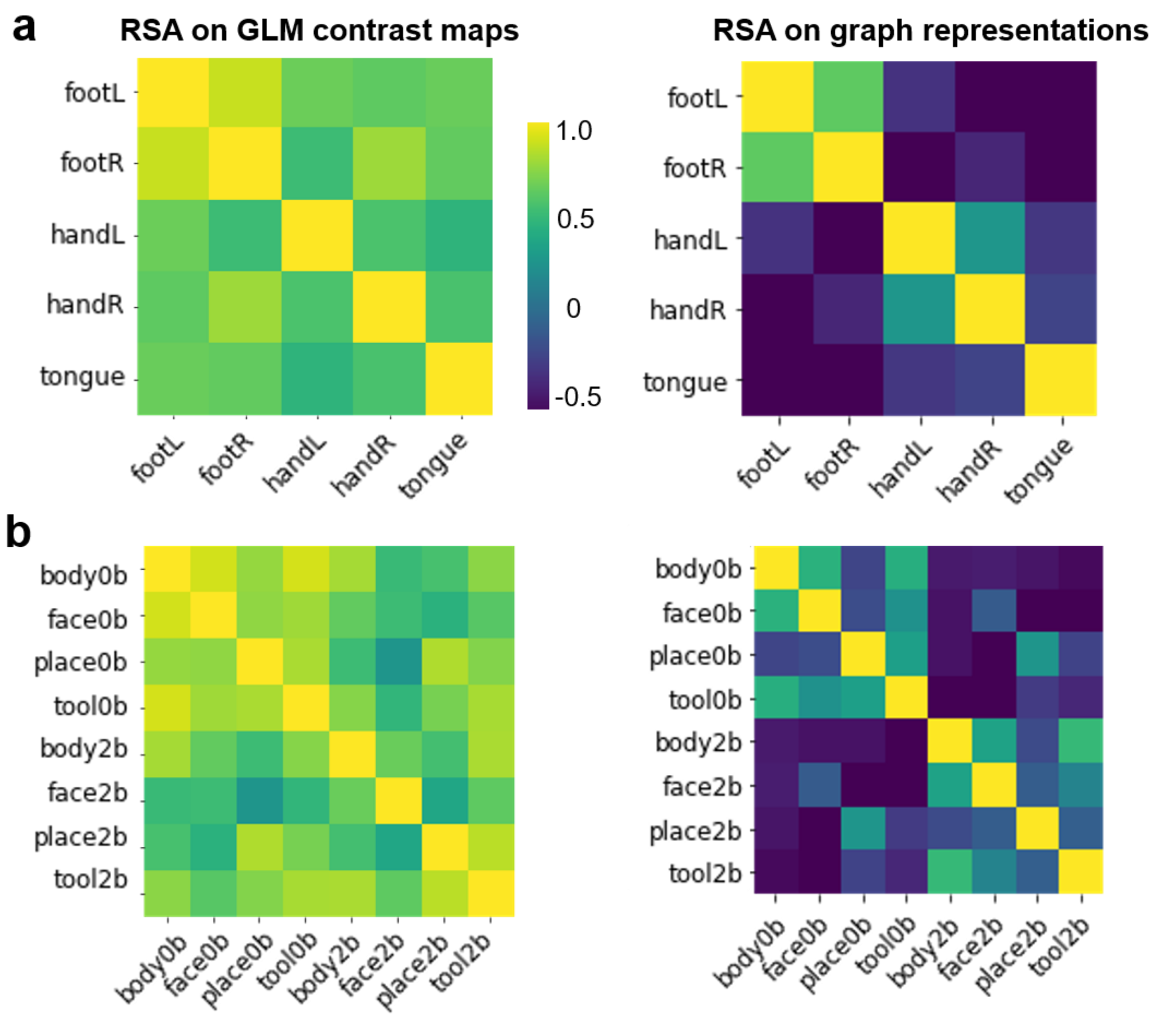


**Fig. 2-S1 | Representational similarity of feature representations using contrast maps and BGNN representations.** The representational similarity of feature representations was evaluated by calculating Pearson correlation coefficients between different MOTOR (**a**) and WM (**b**) tasks. Two types of features were used in this analysis, including contrast maps derived from classical GLM analysis (1st column) and graph representations learned in the BGNN decoding model (2nd column). Compared to the GLM contrast maps, BGNN representations showed higher distinction between movement types (**a**), and 0back *vs* 2back WM tasks (**b**).

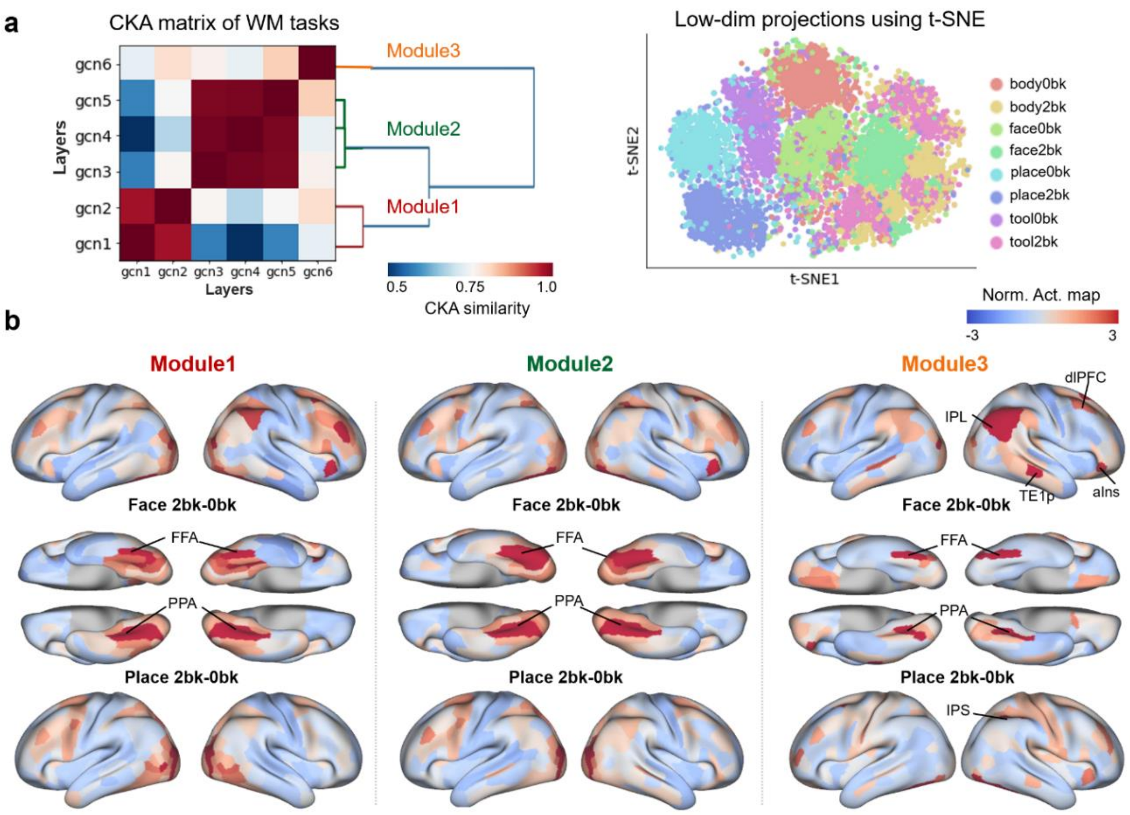


**Fig. 2-S2 | Representational power of the *ChebNet-K1* model as measured by the double-dissociation between memory load and image category.** The *ChebNet-K1* model showed weak power in detecting differential brain mechanisms of representational abstraction for remembering faces vs places, i.e. the contrast of 2back vs 0back tasks on faces vs places. **a),** Similarity of representations between BGNN layers was calculated using CKA with a linear kernel. Three modules were identified in the *ChebNet-K1* model by applying hierarchical clustering to the CKA matrix: *Module1* (gcn1 to gcn2), *Module2* (gcn3 to gcn5), and *Module3* (gcn6). The representations of the last BGNN layer were projected onto a 2-dimensional space using t-SNE. Compared to the *ChebNet-K5* model, the K=1 model showed weaker distinctions among 2back tasks with different visual stimuli. **b),** Differential abstract representational states for remembering faces vs places, i.e. double-dissociation between faces vs places and 2back vs 0back contrasts. The three modules showed weak power in detecting representations for the memory maintenance, i.e. 2back vs 0back contrasts, but captured the visual representations in the ventral stream, i.e. recognition of faces vs places. Specifically, a small portion of the frontoparietal network regions, including dlPFC and IPS, were detected in the 2back vs 0back contrasts on faces but not for places. More distinctions were identified in the ventral visual stream, including FFA for faces and PPA for places. aIns: anterior insula; dlPFC: dorsolateral prefrontal cortex; IPS: intraparietal sulcus; IPL: inferior parietal lobe; FFA: fusiform face area; PPA: parahippocampal place area.

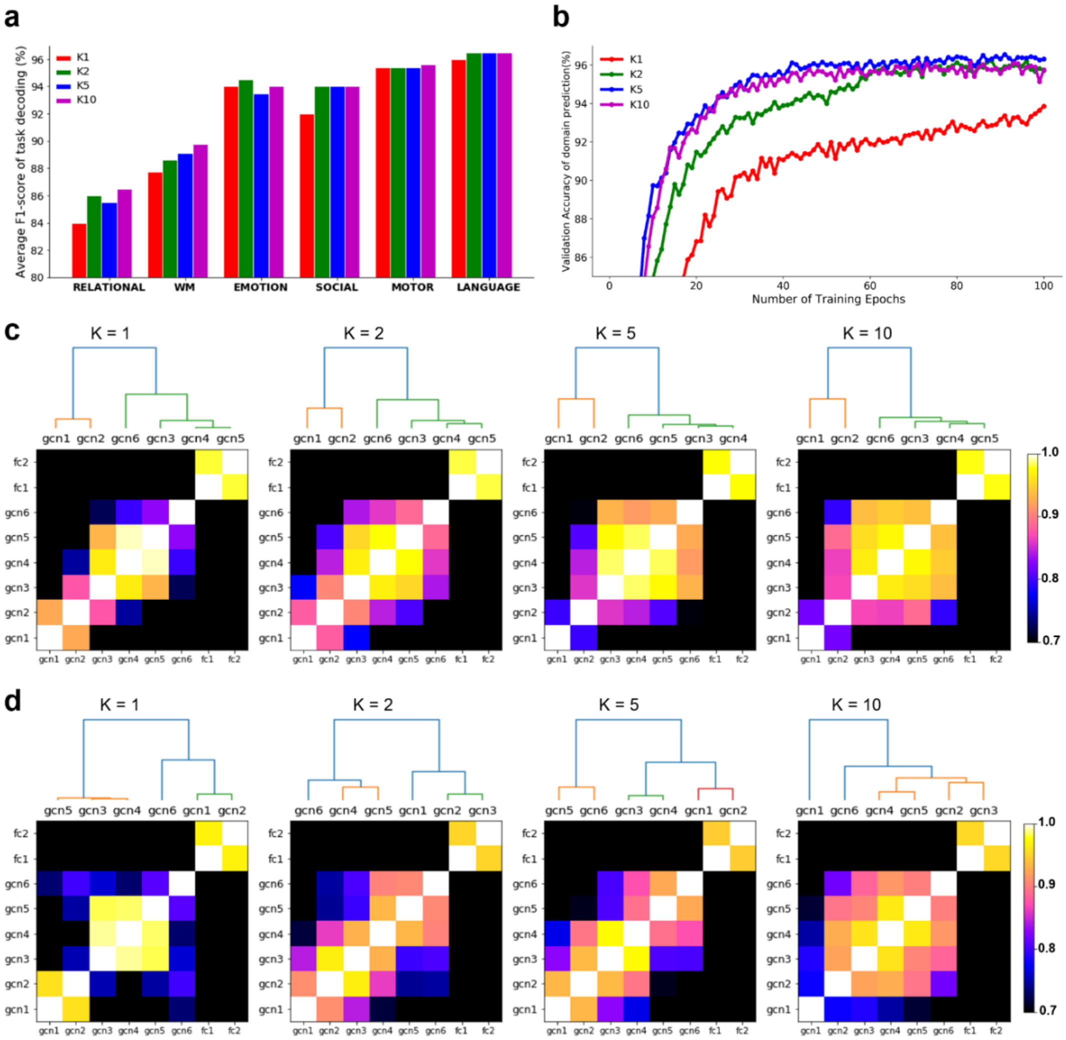


**Fig. 2-S3 | The effect of K-order on brain decoding and hierarchical organization of BGNN on Motor and WM tasks.** The effect of *K*-order on brain decoding was investigated by spanning over the list of [0,1,2,5,10]. The decoding performance on K=0 was not shown in this plot due to its low overall performance (decoding accuracy = 83.76%, 84.21%, 83.51% on training, validation and test sets). High-order decoding models achieved higher decoding accuracy along with faster convergence speeds during model training (**b**). Significant improvements were detected between K=1 (information integration within the same network) and K>1 (transmission of brain activity among inter-connected brain networks). We detected different levels of sensitivity to the choice of *K*-order among a variety of cognitive domains (**a**). The effect of *K*-order on each cognitive domain was estimated by averaging the F1-score on the test set. Among which, the Motor tasks showed stable decoding performance when increasing K while the decoding of WM tasks gradually improved as increasing *K*. Correspondingly, a stable two-level organization among BGNN layers was revealed for the Motor tasks when increasing K (**c**). For the Working-memory task, the decoding performance gradually improved as increasing K and the organizational structure started with an unstable bipartition and gradually evolved into a tripartite organization among BGNN layers (**c**). The similarity of representations among BGNN layers was calculated using CKA with a linear kernel. The hierarchical clustering was then applied to the distance matrix (dis = 1-cka) using Ward's linkage and revealed the organizational principles among BGNN layers.

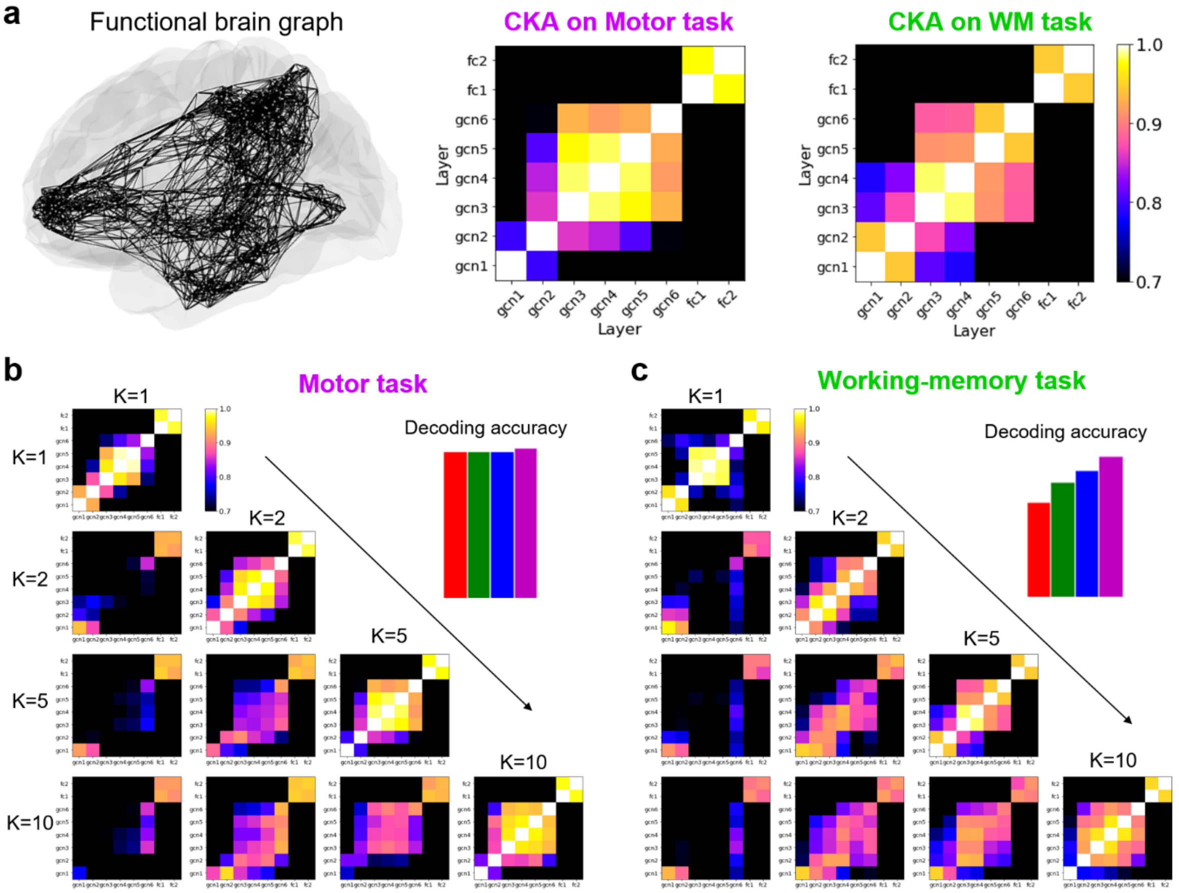


**Fig.2-S4 | Similarity analysis of graph representations derived from different BGNN models on Motor and WM tasks.** The choice of K-order not only affects the decoding performance but also changes the organizational structure of the decoding model. To quantify this effect, we evaluated the similarity of graph representations between BGNN layers as well as between different models using CKA. **a)**, Similarity of BGNN representations for the Motor and WM tasks. The graph convolution was applied on the functional-graph derived from the resting-state functional connectivity. **b)** and **c)**, Similarity of BGNN representations at different K-orders. For the Motor task (**b**), the BGNN model reached the best decoding performance at $K = 1$, and learned similar low-level and high-level representations at different K, indicating that *ChebNet-K1* was enough to capture the localized brain activity during Motor tasks. For the Working-memory task (**c**), the BGNN model showed high sensitivity to the choice of *K*-order and achieved the best decoding accuracy when $K = 10$. The BGNN models learned similar low-level representations at different K, but captured different high-level representations, corresponding to the distinguishable features between 2back and 0back tasks in high-order models, indicating the important role of between-network integration in WM tasks.

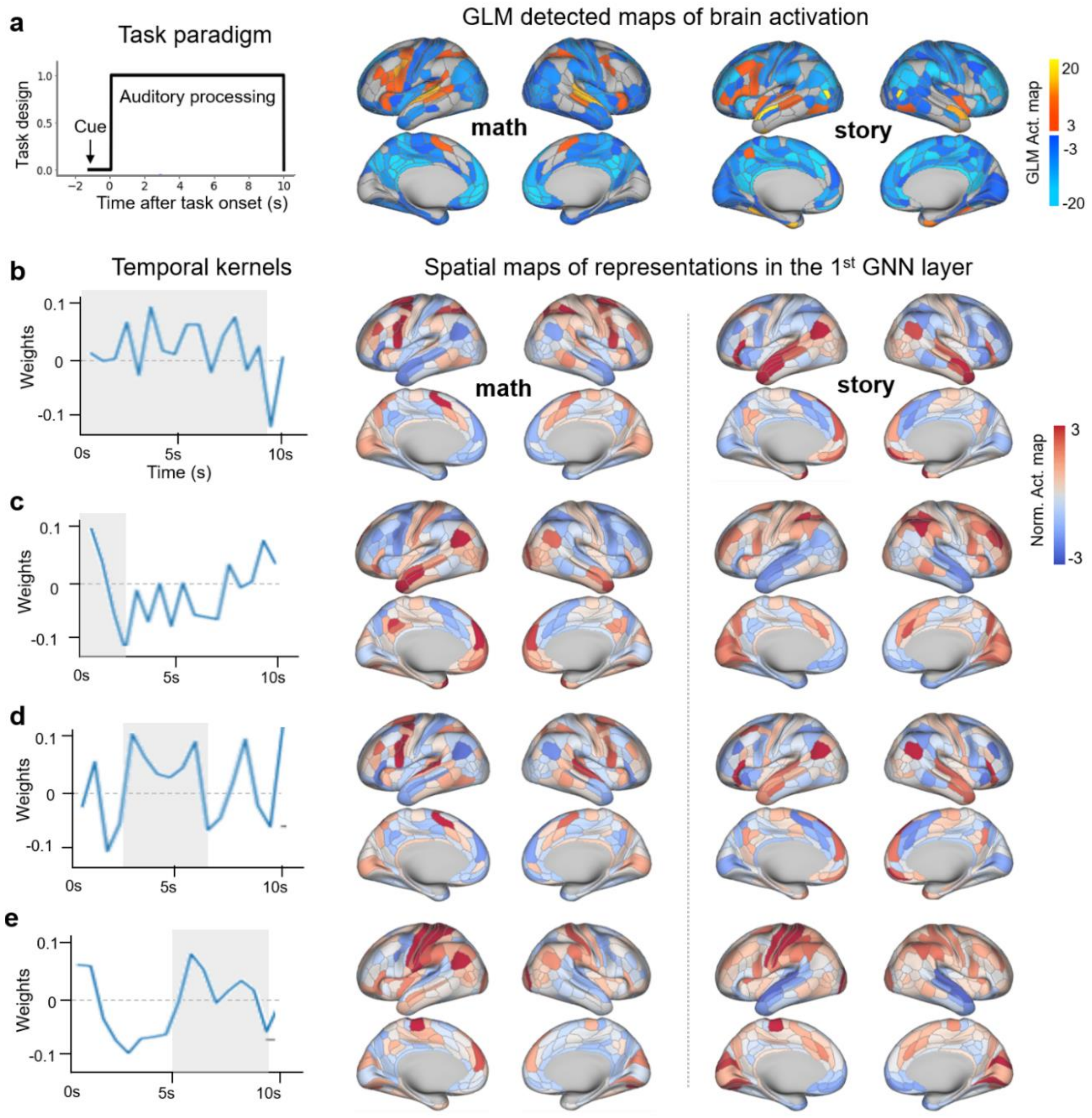


**Fig. 4-S1. Spatiotemporal decomposition of brain responses during the Language tasks.** We use the Language tasks as examples to illustrate that BGNN captures various hemodynamic responses in the temporal domain and brain activations in the spatial domain. **a)**, The task paradigm of each trial and the corresponding brain activation maps detected by the classical GLM analysis. The contrast maps of "math" and "story" conditions were calculated on HCPS500 subjects, downloaded from the neurovault (https://neurovault.org/collections/457/). Each "math" trial starts with ~1s cue, and then presents a ~6s audio of mathematical operations, followed by 3s question and 3s response of button press. The "story" trial comes with longer duration for auditory processing (~20s) and shorter time of question. **b-e)**, Various temporal convolutional kernels were learned at the first BGNN layer ($1^{st}$ column), capturing brain responses at different stages of cognitive processes, for instance, the *cue phase* (**c**), *auditory processing* (**d**) and *button pressing* (**e**), as well as the entire block (**b**). The corresponding "activation map" at each stage and for each task condition, e.g. story ($2^{nd}$ column) and math ($3^{rd}$ column), were also captured. We found that 1) the visual cortex and prefrontal regions were activated during the *cue phase*; 2) the auditory cortex and other task-related regions were activated for the stage of a*uditory processing*, e.g. frontoparietal regions for "math" and perisylvian language areas for "story"; 3) the motor and somatosensory cortex was activated for the stage of *button pressing.* This analysis uncovers a functional gradient in the spatiotemporal organization of task-evoked brain activity.

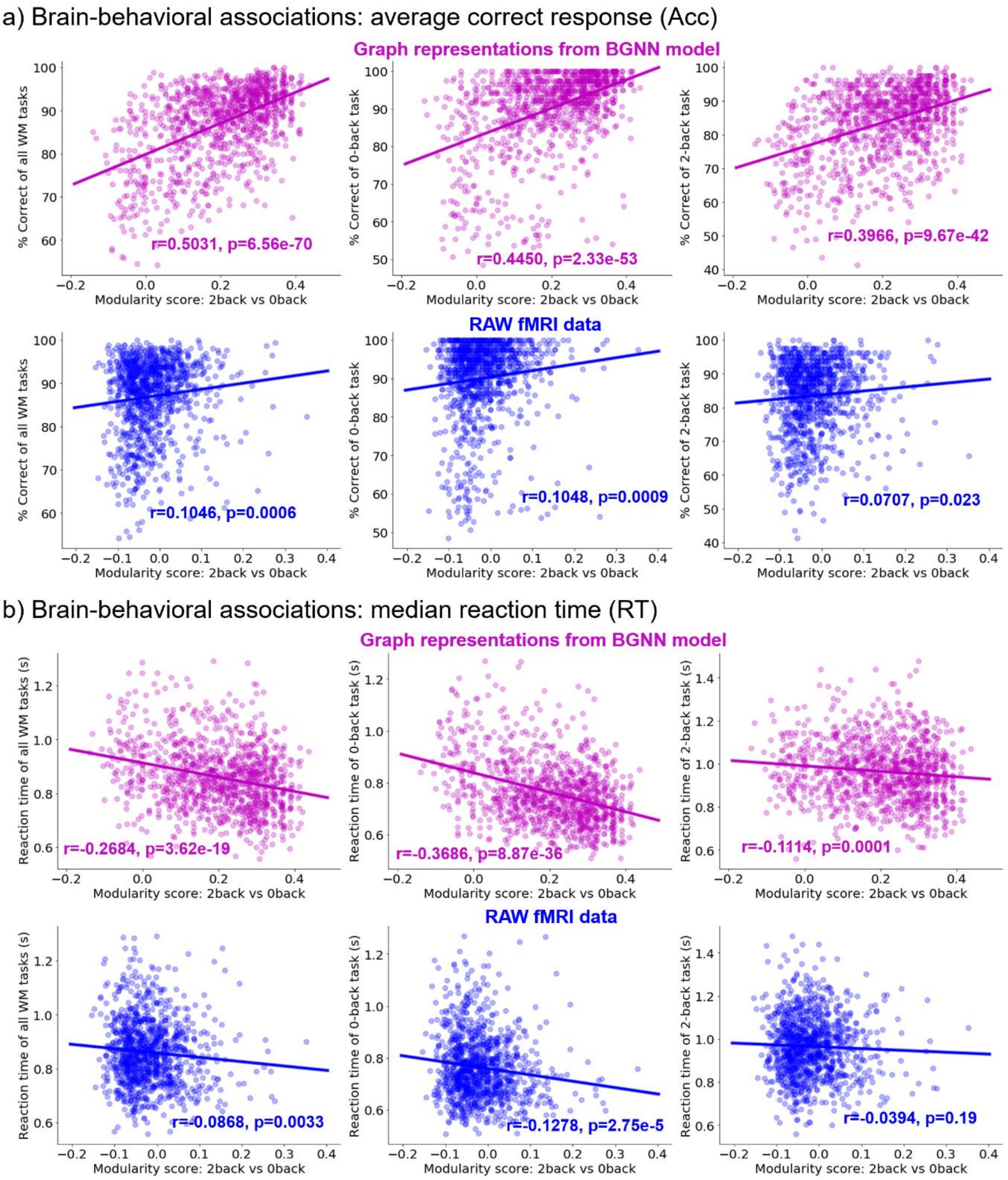


**Fig. 5-S1 | Modularity scores in the state-transition graph significantly correlated with correct responses (Acc) and reaction time (RT) during Working-memory tasks.** The modularity score was calculated based on the state-transition graph of BGNN representations or raw fMRI data from each subject. Specifically, we first constructed a kNN graph (k=5) from the t-SNE projections of learned BGNN representations from each subject. A modularity score was then evaluated on the individual kNN graph with the partition provided by task conditions (e.g. 0back vs 2back). The resulting modularity scores were used to measure the task segregation effect in the learned representations, and were correlated with participant's behavioral performance. **a**), We found significant correlations between the modularity scores of BGNN representations and averaged correct responses (Acc) for all working-memory tasks (1$^{st}$ panel), 0back tasks (2$^{nd}$ panel) and 2back tasks (3$^{rd}$ panel). Much weaker associations were detected in the raw fMRI data. **b**), We found significant correlations between the modularity scores of BGNN representations and median reaction time (RT) for all working-memory task (1$^{st}$ panel), 0back tasks (2$^{nd}$ panel) and 2back tasks (3$^{rd}$ panel). Much weaker associations were detected in the raw fMRI data. The purple lines indicated the linear regression models between the state-transition graph of BGNN representations and behavioral performance. The blue lines indicated the linear regression models between the state-transition graph of raw fMRI data and behavioral performance. The analysis was done among all subjects from HCP S1200 release, with complete records of behavioral and imaging data for working memory tasks (N=1074).

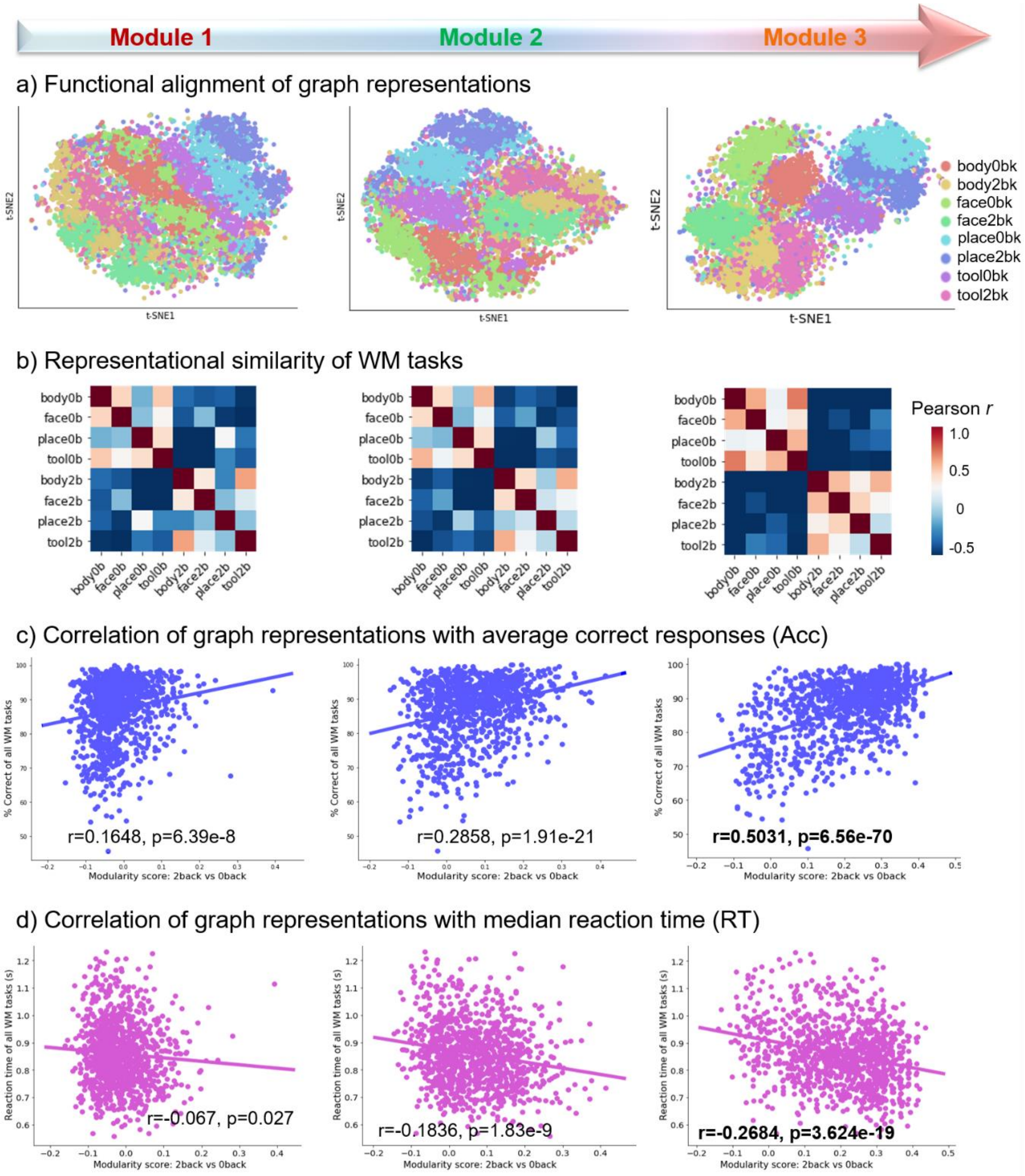


**Fig. 5-S2 | Functional alignment and brain-behavioral associations along the representational hierarchy.** Three modules were identified from the representations of different BGNN layers, i.e. *Module1*, *Module2* and *Module3*. **a**), Functional alignment of BGNN representations. The representations of each module were projected onto a 2-dimensional space by using t-SNE. **b**), The representational similarity of WM tasks was evaluated by calculating Pearson correlation coefficients in the BGNN representations at different representational levels. Both t-SNE projections and representational similarity analysis demonstrated a gradually enhanced effect of *2bk-0bk* along the representational hierarchy, i.e. from low-level representations in *Module1* to high-level representations in *Module3*. **c**) and **d**), Brain-behavioral associations between learned BGNN representations and participants' behavioral performance. The modularity score of *2bk-0bk* in the BGNN representations was significantly correlated with participants' behavioral scores, including both average correct responses (Acc, in **b**) and median reaction time (RT, in **c**). As going deeper along the representational hierarchy, both functional alignment of BGNN representations and the brain-behavior associations were gradually enhanced, indicating a gradual progression towards behaviorally relevant representations.

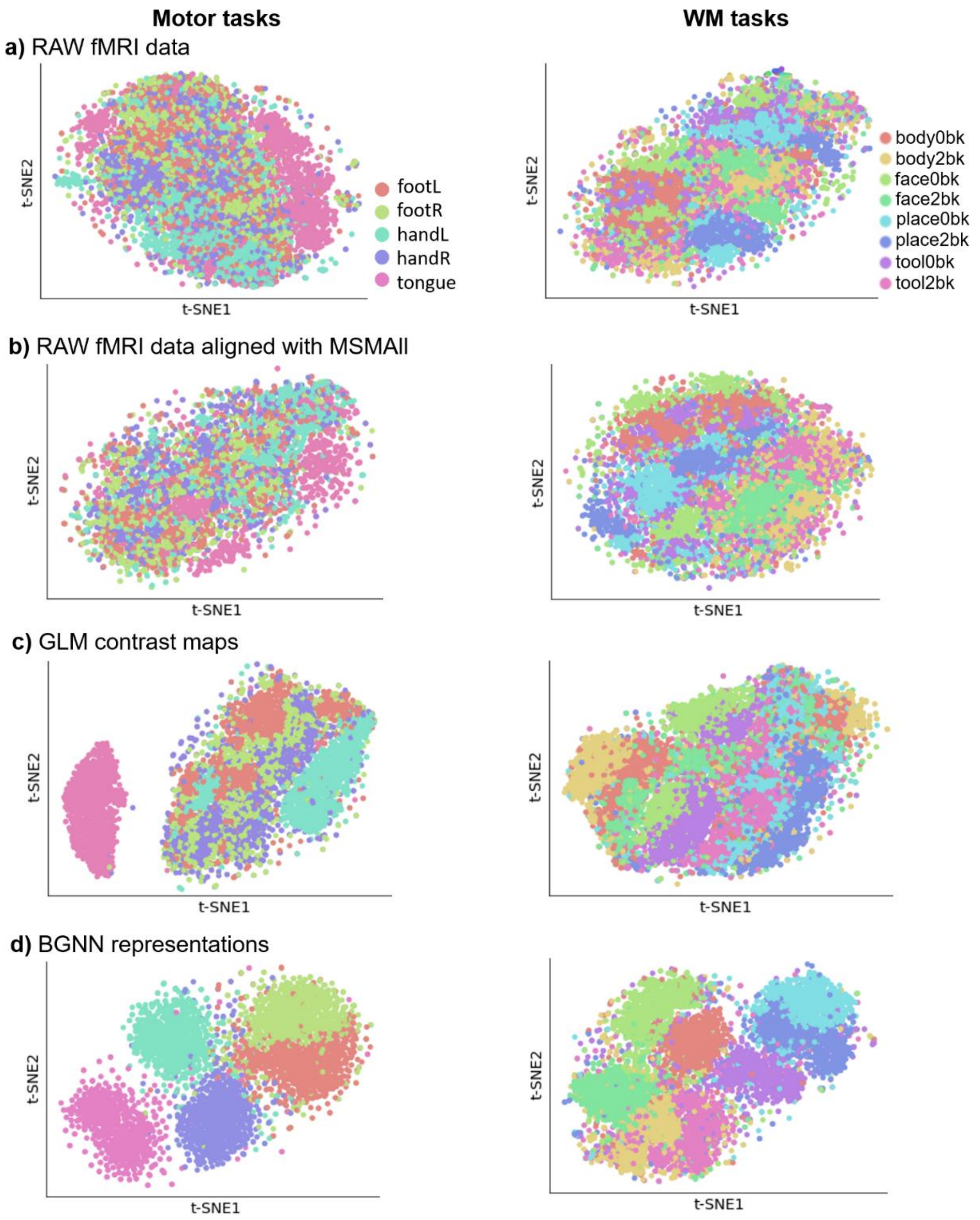


**Fig. 5-S3 | Projection of brain responses using different data registration approaches.** We extracted different types of representations from task fMRI data, including raw fMRI data with and without the "MSMAll" alignment, contrast maps derived from the classical GLM analysis, as well as BGNN representations. All representations were first mapped onto the same brain parcellation, i.e. Glasser's atlas (Glasser et al., 2016) and then projected onto a 2-dimensional space using t-SNE. For both Motor (1st column) and WM tasks (2nd column), BGNN representations showed the best inter-subject alignment of brain responses, with much higher segregation effect than raw fMRI data using different data registration (Q=0.25, 0.28, 0.53, 0.68 for body movements in Motor tasks, Q=0.06, 0.10, 0.16, 0.38 for memory load in WM tasks, respectively for raw fMRI data using “MSMSulc” and “MSMAll” approaches, GLM contrast maps and BGNN representations). Each dot in the plot represents a single trial of a specific task condition (in different colors). The Motor task data includes five types of body movements. The WM task data includes eight task conditions.

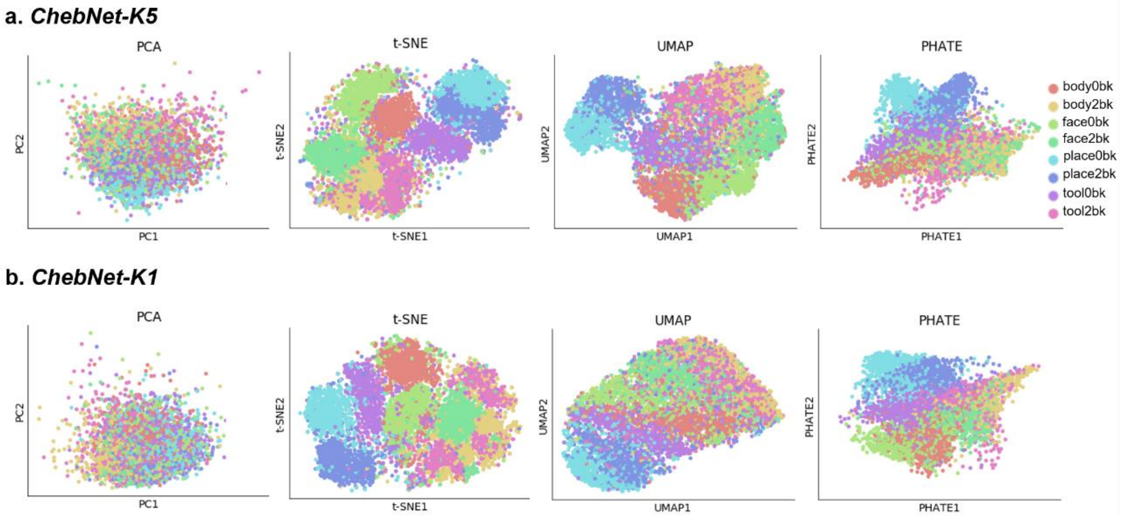


**Fig. 5-S4 | Projection of BGNN representations using different dimension reduction approaches.** BGNN representations were mapped onto a 2-dimensional space by using different dimension reduction techniques, including PCA (1$^{st}$ column), t-SNE (2$^{nd}$ column) (Maaten and Hinton, 2008), UMAP (3$^{rd}$ column) (McInnes et al., 2018), and PHATE (4$^{th}$ column) (Moon et al., 2019). Among which, the best visualization was provided by t-SNE. Here, we evaluated two different decoding models, including *ChebNet-K5* (**a**) and *ChebNet-K1* (**b**). Compared to the low-order model, *ChebNet-K5* model showed higher distinctions among eight WM task conditions, especially between the 2back vs 0back tasks.

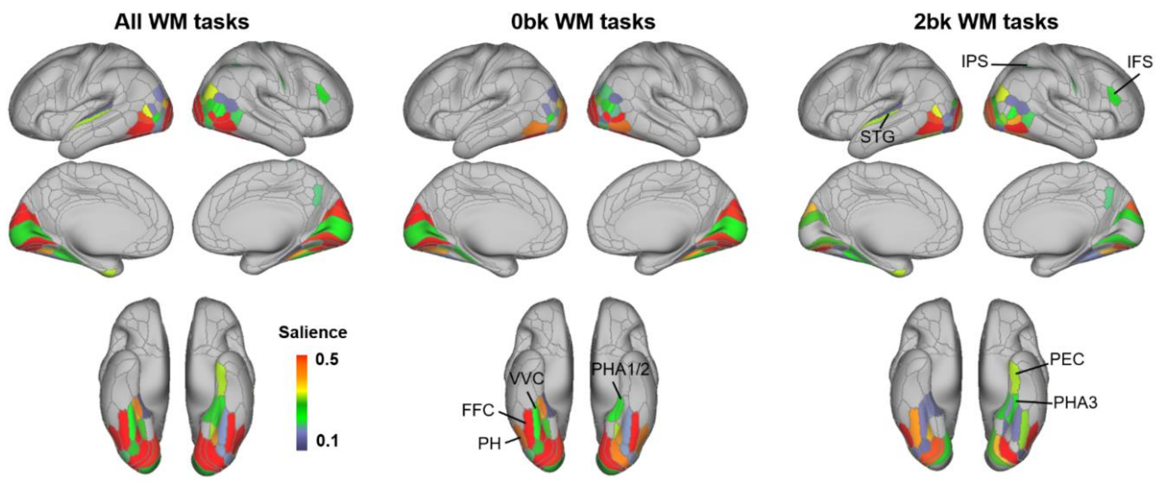


**Fig. 6-S1 | Saliency maps for the decoding of WM tasks.** We generated the salient features that showed high contributions to the classification of WM tasks by using the guided backpropagation approach. The model detected biologically meaningful and category-specific salient features on 0back and 2back WM tasks. For 0back tasks, the model identified salient regions in the visual cortex and ventral visual stream, including V1-V3, PH, FFC and PHA. For 2back tasks, the model detected salient features in the frontoparietal network regions, including IPS and MFG, and the temporal regions, e.g. superior temporal gyrus (STG) and Perirhinal Cortex (PRC). Note that only brain regions with a high saliency (saliency values>0.1, full range of saliency is (0,1)) and a significant 'task condition' effect (p<0.001) were shown in the brain maps.

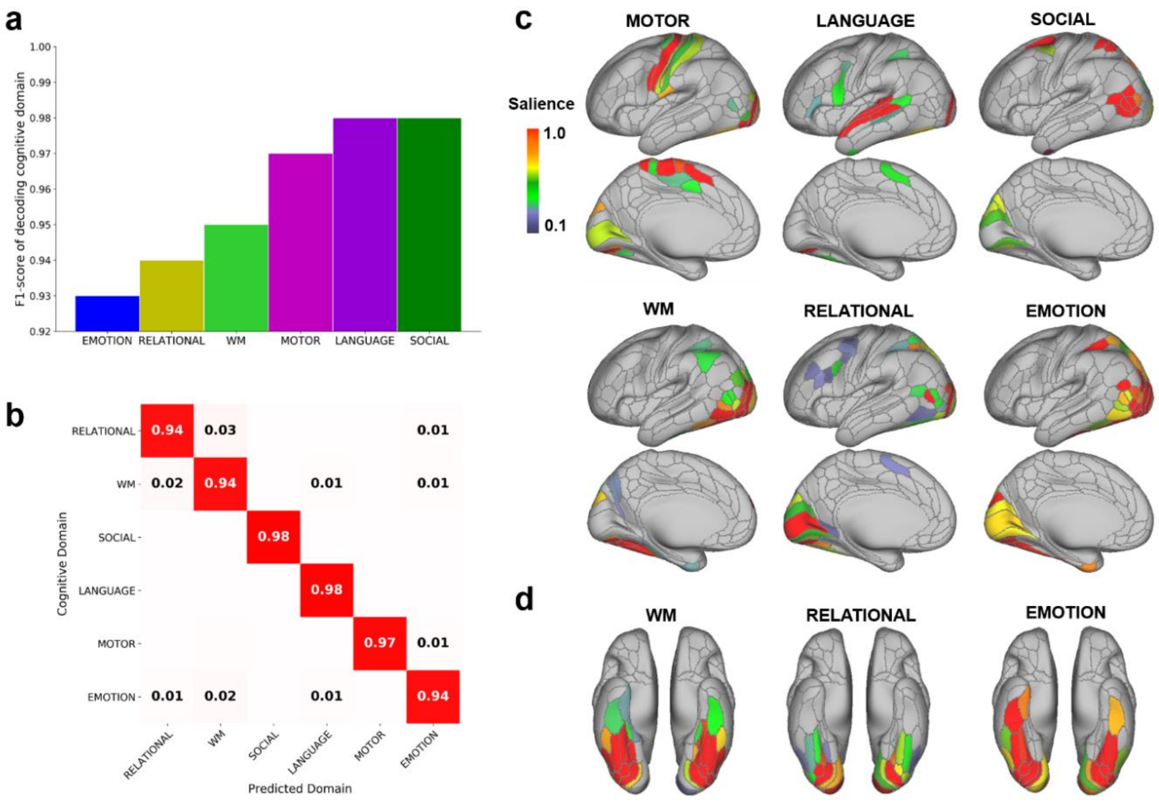


**Fig. 6-S2 | Decoding of six cognitive domains and their saliency maps.** The decoding model predicted the six cognitive domains from each 10s of fMRI responses and achieved an average test accuracy of 96%. **a)**, The F1-score of decoding on each cognitive domain. **b),** The confusion matrix of cross-domain classification. **c),** The saliency map of each cognitive domain in the lateral and medial surfaces. The saliency value indicates the importance of input features that highly contribute to the prediction output of the decoding model. We observed different sets of salient brain regions for each of six cognitive domains. **d)**, The salient brain regions in the ventral temporal cortex. Among the six domains, three tasks involve the processing of visual stimuli and pattern matching in the fMRI paradigm, i.e. Working-Memory (WM), Relational Processing (RELATIONAL) and Emotion (EMOTION). Correspondingly, we detected different sets of salient features in the ventral visual stream for the three tasks.

### Supplementary Tables

**Table S1. Scanning parameters and experimental designs of HCP task-fMRI dataset.**

The entire dataset includes in total of 14,895 functional runs across the six cognitive domains with variable lengths of task duration and different number of task conditions for each domain.

| Task Domains | #Subj | #Runs | #Volumes per run | #Trials per run | #Cond. | Min dura. per block (sec) |
|---|---|---|---|---|---|---|
| Working memory | 1085 | 2 | 405 | 8 | 8 | 25 |
| Motor | 1083 | 2 | 284 | 10 | 5 | 12 |
| Language | 1051 | 2 | 316 | 8 | 2 | 10 |
| Social Cognition | 1051 | 2 | 274 | 5 | 2 | 23 |
| Relational processing | 1043 | 2 | 232 | 6 | 2 | 16 |
| Emotion | 1047 | 2 | 176 | 6 | 2 | 18 |

**Table S2. Behavioral differences for the recognition of face vs place images in WM tasks.** We observed significant differences in behavioral performance among different types of visual stimuli. Specifically, the recognition of face images achieved higher accuracy and faster responses than place images, on both **0back** and **2back** conditions, as well as smaller decays in both measures due to memory load (**2back-0back** condition).

| **WM tasks** | | **Correct response (%)** | | **Reaction time (ms)** | |
|---|---|---|---|---|---|
| **Contrasts** | **Cond.** | **T-score** | **p-value** | **T-score** | **p-value** |
| *Face vs Places* | 0back | 7.76 | 1.84e-14 | -2.38 | 0.017 |
| *Face vs Places* | 2back | 12.22 | 2.86e-32 | -9.90 | 3.68e-22 |
| *Face vs Places* | **2back-0back** | **3.21** | **0.0013** | **-5.97** | **3.16e-9** |

**Table S3: Heritability analysis of brain responses and behavioral scores.**

Heritability estimates were conducted for both representations of brain responses and behavioral performance in-scanner, associated with WM tasks, after controlling for confounding effects of age, gender, handedness and head motion. The average accuracy (Acc) and reaction time (RT) showed high heritability estimates of additive genetic effects. For BGNN representations and raw fMRI signals, the high-dimensional data was first projected onto a 2-dimensional space using t-SNE and then the task segregation effect was estimated based on individual state-transition graph (see Method section). Significant heritability estimates were also detected in the BGNN representations but not in raw fMRI signals.

| **Traits** | **$h^2$** | **SE** | **p-value** | **FDR corrected p-value** | **Covariance Explained** |
|---|---|---|---|---|---|
| *Neural representations of brain responses* | | | | | |
| **BGNN representations-functional graph** | **0.3597** | 0.0522 | **1.04E-11** | 2.08E-11 | 0.0025 |
| **BGNN representations-structural graph** | **0.3493** | 0.0529 | **5.94E-11** | 1.19E-10 | 0.0045 |
| Raw fMRI signals | 0.1148 | 0.0567 | 0.0186 | 0.0186 | 0.0066 |
| *Behavioral measures of WM tasks* | | | | | |
| WM_Task_Acc | 0.5624 | 0.0435 | 8.56E-27 | 3.42E-26 | 0.0434 |
| WM_Task_2bk_Acc | 0.5887 | 0.0425 | 6.97E-29 | 5.58E-28 | 0.0446 |
| WM_Task_0bk_Acc | 0.3215 | 0.0564 | 6.21E-09 | 8.29E-09 | 0.0222 |
| WM_Task_RT | 0.4118 | 0.0560 | 4.01E-13 | 8.01E-13 | 0.0105 |
| WM_Task_2bk_RT | 0.4534 | 0.0556 | 5.40E-15 | 1.44E-14 | 0.0094 |
| WM_Task_0bk_RT | 0.3294 | 0.0583 | 6.06E-09 | 8.29E-09 | 0.0085 |

### References

Bressler, S.L., Menon, V., 2010. Large-scale brain networks in cognition: emerging methods and principles. Trends in Cognitive Sciences 14, 277–290. https://doi.org/10.1016/j.tics.2010.04.004

Cohen, J.R., D'Esposito, M., 2016. The Segregation and Integration of Distinct Brain Networks and Their Relationship to Cognition. J. Neurosci. 36, 12083–12094. https://doi.org/10.1523/JNEUROSCI.2965-15.2016

Glasser, M.F., Coalson, T.S., Robinson, E.C., Hacker, C.D., Harwell, J., Yacoub, E., Ugurbil, K., Andersson, J., Beckmann, C.F., Jenkinson, M., Smith, S.M., Van Essen, D.C., 2016. A multi-modal parcellation of human cerebral cortex. Nature 536, 171–178. https://doi.org/10.1038/nature18933

Maaten, L. van der, Hinton, G., 2008. Visualizing Data using t-SNE. Journal of Machine Learning Research 9, 2579–2605.

McInnes, L., Healy, J., Saul, N., Großberger, L., 2018. UMAP: Uniform Manifold Approximation and Projection. Journal of Open Source Software 3, 861. https://doi.org/10.21105/joss.00861

Moon, K.R., van Dijk, D., Wang, Z., Gigante, S., Burkhardt, D.B., Chen, W.S., Yim, K., Elzen, A. van den, Hirn, M.J., Coifman, R.R., Ivanova, N.B., Wolf, G., Krishnaswamy, S., 2019. Visualizing structure and transitions in high-dimensional biological data. Nat Biotechnol 37, 1482–1492. https://doi.org/10.1038/s41587-019-0336-3

Springenberg, J.T., Dosovitskiy, A., Brox, T., Riedmiller, M., 2014. Striving for Simplicity: The All Convolutional Net.
